# Extraordinary genome instability and widespread chromosome rearrangements during vegetative growth

**DOI:** 10.1101/304915

**Authors:** Mareike Möller, Michael Habig, Michael Freitag, Eva H. Stukenbrock

**Affiliations:** Environmental Genomics, Christian-Albrechts University, Am Botanischen Garten 1-9, D-24118 Kiel, Germany; Max Planck Institute for Evolutionary Biology, August-Thienemann-Str. 2, D-24306 Plön, Germany; Department of Biochemistry and Biophysics, Oregon State University, Corvallis, OR 97331-16 7305, United States of America

## Abstract

The haploid genome of the pathogenic fungus *Zymoseptoria tritici* is contained on “core” and “accessory” chromosomes. While 13 core chromosomes are found in all strains, as many as eight accessory chromosomes show presence/absence variation and rearrangements among field isolates. We investigated chromosome stability using experimental evolution, karyotyping and genome sequencing. We report extremely high and variable rates of accessory chromosome loss during mitotic propagation *in vitro* and *in planta*. Spontaneous chromosome loss was observed in 2 to >50 % of cells during four weeks of incubation. Similar rates of chromosome loss in the closely related *Z. ardabiliae* suggest that this extreme chromosome dynamic is a conserved phenomenon in the genus. Elevating the incubation temperature greatly increases instability of accessory and even core chromosomes, causing severe rearrangements involving telomere fusion and chromosome breakage. Chromosome losses do not impact the fitness of *Z. tritici in vitro*, but some lead to increased virulence suggesting an adaptive role of this extraordinary chromosome instability.

## Introduction

Pathogenic fungi pose global threats to agriculture, human, and animal health. Pathogens infecting plants and animals have been shown to rapidly adapt to changing environments, including industrial agriculture and medical treatments, usually in response to adaptive pressure caused by the widespread use of fungicides and pharmaceuticals (Selmecki et al. 2010; Bennett et al. 2014). The genomes of many prominent fungal plant pathogens exhibit high levels of structural variation including isolate– or lineage-specific regions often characterized by high repeat contents, and accessory chromosomes varying in frequency among individual isolates (De Jonge et al. 2013; Persoons et al. 2014; Plissonneau & Stürchler 2016). Structural variation is often associated with meiotic recombination, but highly variable regions are also found in asexually and mostly asexually reproducing species (Faino et al. 2016; Chuma et al. 2011; Wang et al. 2006). In several plant pathogenic fungi, determinants of virulence, so called effector proteins, are located in dynamic regions of the genome (Coleman et al. 2009; Ma et al. 2010; V. Miao et al. 1991). Accessory chromosomes often carry genes that encode virulence determinants and are linked to pathogenicity. Therefore, the absence of a particular chromosome in some species results in avirulent phenotypes (Tsuge et al. 2016; Vlaardingerbroek et al. 2016; Ma et al. 2010; van Dam et al. 2018). Little is known about the mechanisms that drive the dynamics of accessory chromosomes and highly variable regions in genomes of eukaryotic pathogens.

The plant pathogenic fungus *Zymoseptoria tritici* causes disease on wheat, especially in Northern Europe and North America, resulting in extensive annual yield loss (Fones & Gurr 2015; Torriani et al. 2015). The *Z. tritici* reference isolate IPO323 contains eight accessory chromosomes with sizes ranging from 0.4 to 1 Mb, accounting for 12% of the entire genome (Goodwin, Ben M’Barek, et al. 2011). Chromosome rearrangements, including complete loss of accessory chromosomes, are a frequent phenomenon in this fungus and have been widely studied as a consequence of meiosis (Wittenberg et al. 2009; Croll et al. 2015; Fouche et al. 2017). Infection experiments with *Z. tritici* strains in which single or multiple accessory chromosomes had been deleted showed that at least some accessory chromosomes encode virulence factors that determine host specificity (Habig et al. 2017). In contrast to the core chromosomes, accessory chromosomes are enriched with transposable elements and have low gene density (Goodwin, M’barek, et al. 2011). Chromatin of accessory chromosomes shows hallmarks of constitutive and facultative heterochromatin, consistent with the observed transcriptional silencing of genes present on these chromosomes (Schotanus et al. 2015; Kellner et al. 2014; Rudd et al. 2015). The centromeres, subtelomeric regions and telomeric repeats of accessory chromosomes are indistinguishable from those of core chromosomes (Schotanus et al. 2015), suggesting that accessory chromosomes contain all the required regions for proper chromosome segregation. The unusual high number of accessory chromosomes, in combination with the extreme variability in chromosome content among field isolates or progeny from controlled crosses, make *Z. tritici* an excellent eukaryotic model to study accessory chromosomes and their dynamics (Mehrabi et al. 2007; Stukenbrock et al. 2010; Goodwin, M’barek, et al. 2011; Wittenberg et al. 2009). Here, we provide evidence for unexpectedly high rates of chromosome loss and changes during asexual propagation, both *in vitro* and *in planta*. We describe the types of structural rearrangements and overall variation in chromosome stability in this important crop pathogen.

## Results

### Accessory chromosomes are lost at a very high rate *in vitro*

To assess the stability of accessory chromosomes in *Z. tritici* we conducted an *in vitro* longterm growth experiment. We used the *Z. tritici* isolate IPO323Δchrl8, derived from the reference strain IPO323 for which there is a completely assembled genome (Goodwin, M’barek, et al. 2011). IPO323Achrl8 lost chromosome 18 during previous *in vitro* propagation, and is here referred to as Zt09 (Kellner et al. 2014).

We propagated fungal cells in liquid culture at 18°C, including eight transfers to fresh medium. After four weeks (representing ^~^80 cell divisions per cell), 576 single strains originating from three replicate cultures were tested for the presence of the seven accessory chromosomes from the Zt09 progenitor by a PCR assay. The presence of marker regions, located close to the centromere, and the right and left telomere repeats of each accessory chromosome, were tested (Table S1). When the screening by PCR suggested absence of an accessory chromosome, we further validated the result by electrophoretic separation of the fungal chromosomes by pulsed field gel electrophoresis (PFGE) (Figure 1 A and B). Thirty eight of the 576 (^~^7 %) tested strains lacked one accessory chromosome. We did not find strains that lacked more than one chromosome, but observe a clear trend where accessory chromosomes 14, 15 and 16 were lost more frequently than smaller accessory chromosomes (Table 1). For example, chromosome 14 was absent in eighteen strains, chromosome 15 in eight and 16 in nine strains. The small chromosomes 20 and 21 were lost only in two and one strain, respectively, whereas chromosomes 17 and 19 were never absent following the *in vitro* propagation (Table 1 and Table S2).

**Figure 1:**
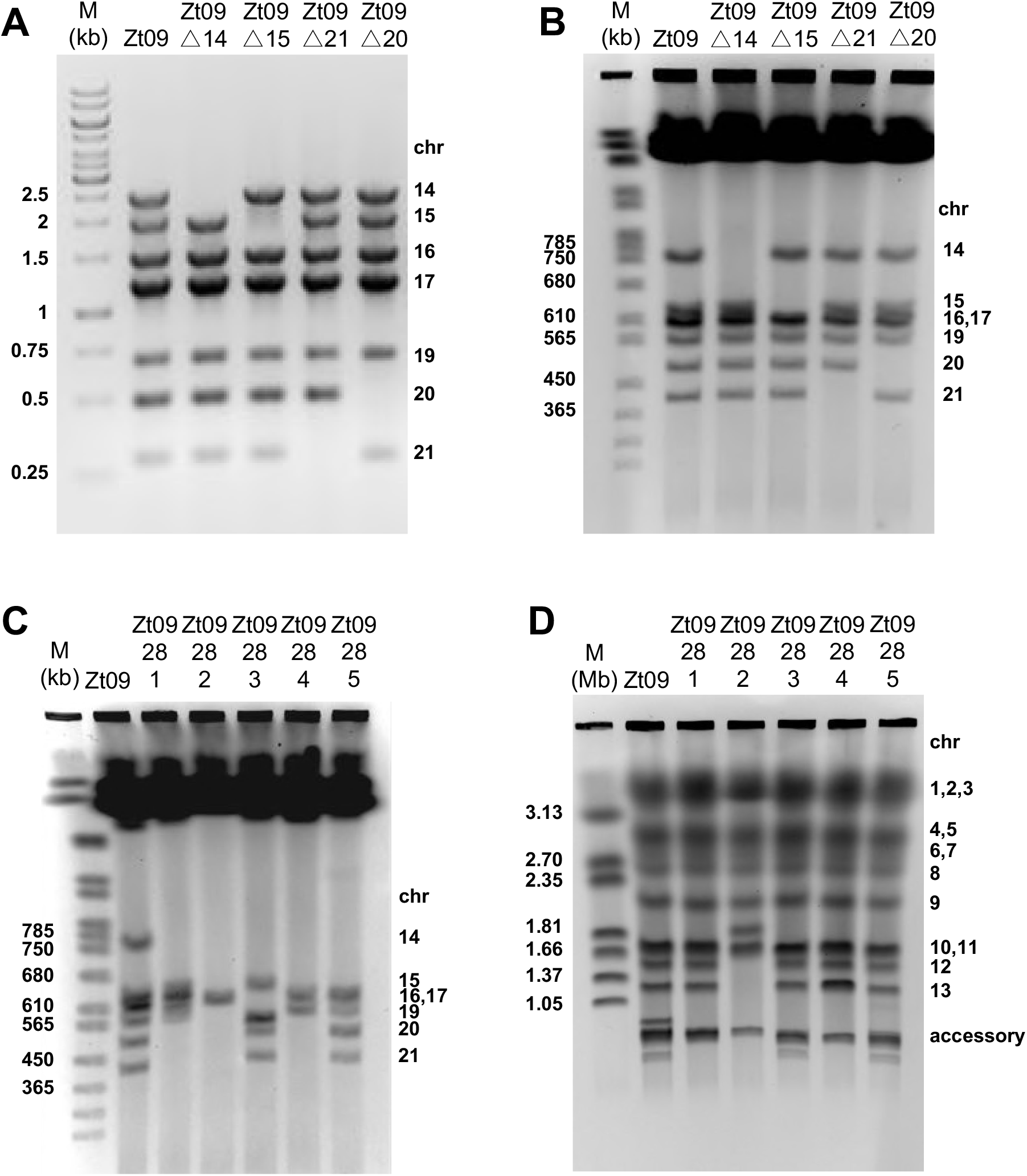
Screening by PCR and pulsed-field gel electrophoresis to identify accessory chromosome-loss strains in *Z. tritici*. (A) Screening by PCR of the progenitor Zt09 strain and evolved strains to assess chromosome losses using primer pairs located close to the centromeric region of the respective accessory chromosome. Multiplex PCR was used to simultaneously screen for the presence or absence of all accessory chromosomes in one strain. Absence of a PCR product indicates the absence of the chromosome; chromosome 18 is not present in Zt09. (B) Pulsed-field gel electrophoresis (PFGE) was conducted to validate the absence of accessory chromosomes in chromosome loss candidates identified by the initial PCR. Here, the separation of small accessory chromosomes of the progenitor strain Zt09 and four chromosome-loss strains is shown. The corresponding chromosome to each band is indicated on the right. The absence of chromosomal bands confirms the loss of the respective chromosome; chromosome 18 is similar in length to 16 and 17 but is absent in Zt09. PFGE of the accessory (C) and mid-size (D) chromosomes of strains originating from the *in vitro* temperature stress experiment confirms multiple accessory chromosome losses, size alterations and chromosome fusions (see also Figures 3 and S1 and Table S3). The bands for chromosomes 3, 12, 13 and 21 are absent in strain 28-2, however, genome sequencing confirms that the chromosomes are still present in the genome (Figure 2C). The absence of bands on the PFGE can be explained by chromosome size changes whereby chromosomes 3 and 13 and 12 and 21 have experienced chromosome fusions (Figure 3 and S1). Note a new chromosome band with a size of ^~^1.8 Mb resulting from the fusion of chromosomes 12 and 21. In the strain 28-3, the accessory chromosomes 17 and 19 are shorter than the reference chromosomes due to chromosome breakage (Figure 2C and Table S3). Strain 28-4 has an additional band of ^~^1.2 Mb, representing the fusion of a duplicated chromosome 17 (Figure S1). All images of stained gels are color-inverted to make differences more obvious.

**Table 1:** Summary of chromosome losses in *Z. tritici* during evolution experiments *in vitro* and *in planta*. Listed are the number of cells that lost an accessory chromosome and which accessory chromosomes were lost. Sizes of the accessory chromosomes are listed next to the chromosome number. Strains with more than one chromosome lost are listed in brackets.

| Chromosome<br>(kb) | Zt09 *in vitro*<br>culture | Zt09 *in vitro*<br>plate | IPO323 *in*<br>*planta* | Zt09 *in vitro*<br>temperature<br>stress |
| --- | --- | --- | --- | --- |
| 14 (773) | 18 | 0 | 1 | 108 (33) |
| 15 (639) | 8 | 0 | 6 | 2 (8) |
| 16 (607) | 9 | 4 | 1 | 0 (9) |
| 17 (584) | 0 | 0 | 0 | 0 |
| 18 (574) | N/A | N/A | 7 | N/A |
| 19 (550) | 0 | 0 | 2 | 2 (2) |
| 20 (472) | 2 | 0 | 0 | 1 (17) |
| 21 (409) | 1 | 1 | 0 | 1 (7) |
| **Total chr loss**<br>**strains** | **38** | **5** | **17** | **114 (34)** |
| **Total strains**<br>**tested** | **576** | **40** | **986** | **188** |

We considered the possibility that natural selection acted on a large population of cells in the liquid *Z. tritici* cultures and thereby contributed to the observed non-random chromosome losses. To test this, we designed a second evolution experiment to propagate individual cell lineages in the absence of selection. For this experiment, 40 strains originating from one common progenitor (Zt09) were cultivated on plates. Every week, randomly selected colonies derived from single cells were transferred to a fresh plate. These repeated strong bottlenecks allowed us to propagate single lineages of *Z. tritici* without an effect of natural selection (Barrick & Lenski 2013). After four weeks of growth (including four transfers to new plates), the 40 evolved strains were tested for the presence of accessory chromosomes as described above. Five out of 40 (^~^13 %) strains, twice as many as in the previous experiment, lacked an accessory chromosome suggesting that selection in a larger population of cells indeed had removed some of the spontaneously occurring chromosome losses. Interestingly, in this experiment we observe the loss of only two different chromosomes: Chromosome 16 was lost in four strains, and chromosome 21 in one strain. Our two *in vitro* experiments suggest that chromosome loss is not entirely random, and that specific chromosomes are lost at a higher rate (Table 1), depending on the *in vitro* growth conditions. The frequency of chromosome losses during asexual growth *in vitro* is extremely high in *Z. tritici* and exceeds previously reported spontaneous chromosome losses in *F. oxysporum* f. sp. *lycopersici* by far (Vlaardingerbroek et al. 2016).

### Accessory chromosomes are unstable *in planta*

The growth of *Z. tritici* in rich medium *in vitro* is highly distinct from growth of the fungus in its natural environment. *Zymoseptoria tritici* is a hemi-biotrophic pathogen infecting the mesophyll of wheat leaves (Goodwin, M’barek, et al. 2011; Ponomarenko et al. 2011), where it forms asexual fruiting bodies, pycnidia, containing presumably clonal conidiospores (pycnidiospores). Pycnidia develop in the sub-stomatal cavities, where the environment and nutrient availability differ from our tested *in vitro* conditions (Haueisen et al. 2017; Rudd et al. 2015). To test whether chromosome loss occurs only *in vitro* or also during the natural lifecycle of *Z. tritici*, we assessed the loss of accessory chromosomes *in planta*. We collected pycnidia from wheat leaves infected with the *Z. tritici* strain IPO323 and tested single pycnidiospores for the presence of accessory chromosomes. A total number of 968 pycnidiospores originating from 42 separate pycnidia were screened by PCR. In total, 17 strains missing an accessory chromosome were identified (^~^1.7%). Chromosomes 15 and 18 were the most frequently lost chromosomes, while chromosomes 17, 20 and 21 were not lost in any of the strains (Table 1 and Table S2). As for the *in vitro* experiments, no strains lacking more than one chromosome were found. Interestingly, chromosome 18, which was found to be lost at high rates in our plant experiments, was shown to be frequently absent in field isolates *of Z. tritici* (Croll et al. 2013; McDonald et al. 2016). In summary, our findings show that chromosome loss in *Z. tritici* not only results from the non-disjunction of homologous chromosomes during meiosis II, as proposed previously (Wittenberg et al. 2009), but also from frequent chromosome losses during asexual growth and mitotic spore formation *in planta*. The observed frequency of chromosome losses *in planta* is not as high as during *in vitro* growth, however, the number of mitotic divisions to develop pycnidia is presumably lower than during four weeks of *in vitro* growth. Therefore, we propose that the reduced number of chromosome losses *in planta* rather reflects the reduced number of cell divisions than possible fitness effects of chromosome losses.

### Chromosome instability is greatly increased during exposure to heat stress

In their natural environments, pathogens are exposed to various kinds of biotic and abiotic stresses. For example, the local environment on the leaf surface can fluctuate severely in temperature and humidity conditions (Zhan & McDonald 2011). We assessed the impact of an increase in temperature on chromosome stability of *Z. tritici*. To this end, we cultivated Zt09 *in vitro* at elevated temperatures, namely an increase of ten degrees Celsius from 18° to 28°C during four weeks of incubation. Subsequent screening by PCR for accessory chromosomes revealed severe chromosome losses in the tested strains. Out of 188 evolved and tested strains, 148 (^~^80 %) were missing at least one accessory chromosome. Chromosome 14 was the most frequently lost chromosome and 34 evolved strains were lacking more than one chromosome. The maximum number of missing chromosomes was six in one strain (Table 1 and Table S2). PFGE revealed that karyotype alterations were not restricted to simple chromosome loss as observed in the *in vitro* experiments at 18°C and the *in planta* experiments. In contrast, we observed frequent size variation of both core and accessory chromosomes as a consequence of chromosome breakage, fusions and duplications (see below) (Figure 1C and 1D, Figure S1, Table S3).

### Genome sequencing of chromosome-loss strains reveals chromosome breakage and fusion, but a low number of SNPs

In total, we sequenced 19 genomes originating from the different *in vitro* and the *in planta* experiment and the respective progenitor strains; all sequencing results have been deposited in the Sequence Read Archive (SRA) under BioProject ID PRJNA428438. Overall, we confirmed the absence of complete accessory chromosomes (Figure 2A), and we found very few additional changes. After filtering to exclude reads of poor quality and coverage (see Materials and Methods, Supplementary Text) we identified a total number of nine SNPs in eight sequenced strains originating from the *in vitro* experiment in liquid culture at 18°C and the *in planta* experiment. Of these, three SNPs were located in coding regions (Table S4). Besides the complete loss of accessory chromosomes and the identified SNPs, we found no other mutations when comparing the progenitor and evolved strains. However, we identified SNPs distinguishing Zt09 from our IP0323 reference strain, and the published genome sequence; most of these SNPs were in non-coding sequences (Table S4). As we show chromosome losses in different strains, we conclude that the few point mutations do not have measurable impact on chromosome loss.

**Figure 2:**
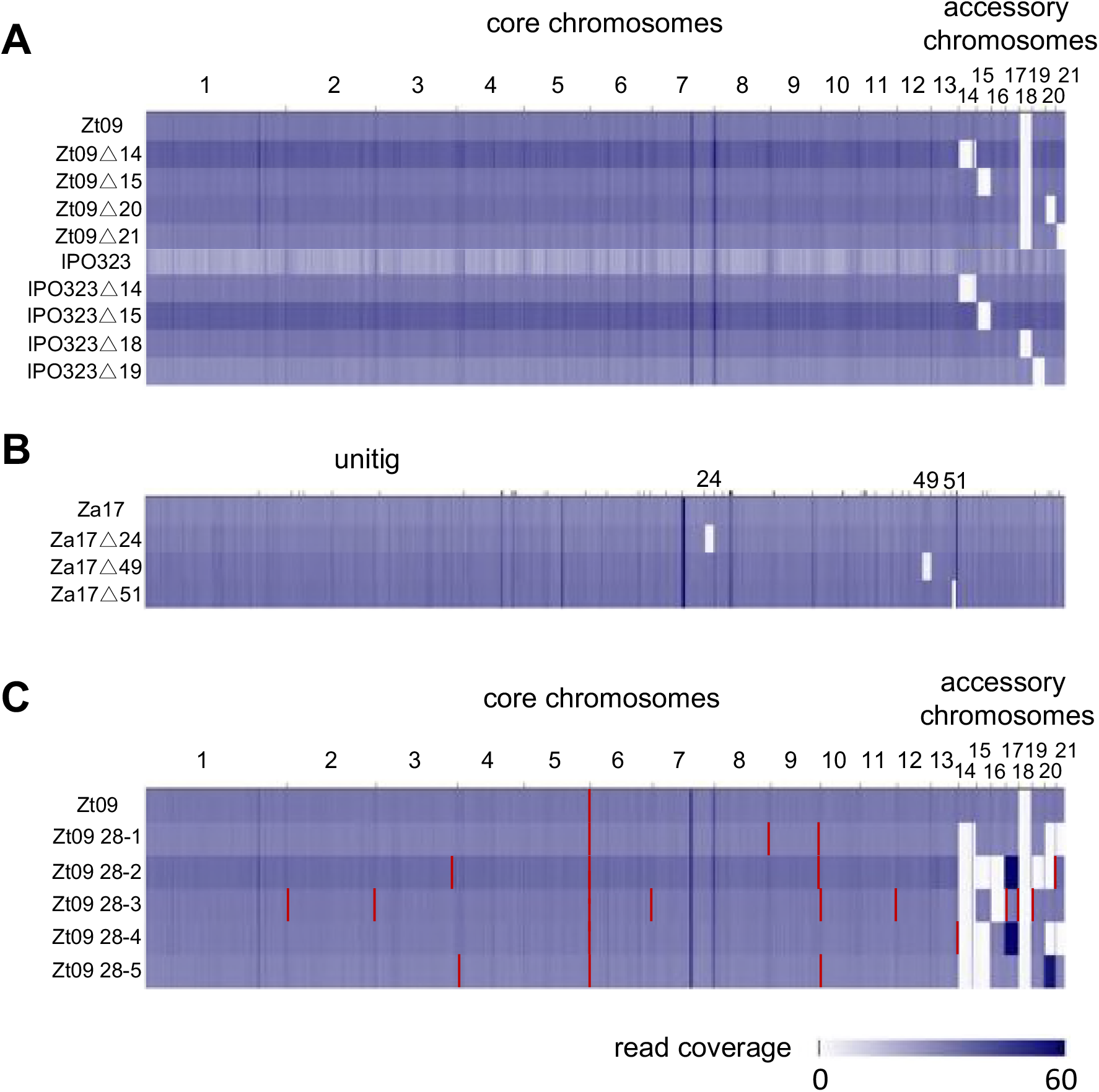
Whole genome sequencing confirms loss of accessory chromosomes in *Z. tritici* and *Z. ardabiliae*. The genomes of chromosome-loss strains derived from A) the *Z. tritici in vitro* evolution experiment and the *in planta* pycnidiospore isolations, and B) the *Z. ardabiliae in vitro* evolution experiment and the respective progenitor strains were sequenced by paired-end Illumina sequencing and mapped to the respective reference isolates, IPO323 (*Z. tritici*) or Za17 (*Z. ardabiliae*). The losses of entire chromosomes (white boxes) were verified for each strain, but few mutations in form of SNPs, INDELs or copy number variation were detected (Table S4). The *Z. tritici* reference genome consists of whole chromosomes, while the *Z. ardabiliae* reference genome is composed of unitigs obtained from the SMRT sequencing assembly (ordered by name of the unitig). C) Genome sequencing of strains derived from the temperature stress experiment revealed, besides verification of accessory chromosome losses, chromosome breaks at the ends of core and accessory chromosomes (red lines) and accessory chromosome duplications. Darker blue shading indicates higher coverage. Most repetitive sequences have a higher coverage than single copy regions resulting in different shades of blue in the coverage graph. The intense dark blue lines indicate high coverage regions on rDNA clusters or repetitive DNA due to underestimation of repeats in the reference assembly (for example, the thin black line in chromosome 7 indicates the location of several rDNA cluster repeats, the only such repeats in the annotated genome sequence, but the expected true number of rDNA repeats is ^~^50).

Whole genome sequencing of five strains evolved at 28°C verified the losses of several accessory chromosomes. Furthermore, comparison of the re-sequenced genomes of the progenitor and the evolved strains revealed substantial size variation resulting from chromosome breakage of core and accessory chromosomes, and duplications of accessory chromosomes (Figure 2C). Chromosome breaks located close to the ends of chromosomes resulted in shortened chromosomes caused by subtelomeric deletions of ^~^0.2-60 kb (Table S3). Detailed analyses of the distribution of discordant paired-end reads mapping to different chromosomes revealed a high number of reads at the chromosome breakpoints with the respective read mates mapping to telomeric repeats (Figure 3). This suggested that most chromosome breaks resulted in shorter chromosomes to which telomeres were added *de novo*. Besides *de novo* telomere formation, we found evidence for fusion of core chromosomes in strain Zt09 28-2, where discordant read mapping indicated fusion of the right arms of chromosomes 3 and 13 (Table S3 and Figure 3). PFGE and Southern blots further showed fusion of chromosomes 12 and 21 in Zt09 28-2 (Figure S1). In a previous study, we reported evidence for the likely fusion of an ancestral accessory chromosome and core chromosome 7 in the reference isolate IPO323 and Zt09 (Schotanus et al. 2015; Kellner et al. 2014). Here, we deduced a fusion of the duplicated chromosome 17 in strain Zt09 28-4 (Figure 1C and D, Figure S1). Similar fusions of the same accessory chromosome 17 following meiosis had been suggested previously, and “breakage-fusion bridge” cycles (McClintock 1938; McClintock 1941) were invoked as a mechanism to form a new accessory chromosome (Croll et al. 2013). In total, in the five analyzed genomes derived from strains grown at 28°C, we identified spontaneous changes: (1) fifteen chromosome breakages (twelve with *de novo* telomere formation and three without evidence for new telomeres), (2) three chromosome fusions, (3) three chromosome duplications, and (4) 16 chromosome losses (Table S3). A key observation from our genome data analyses is the frequent involvement of *de novo* telomere formation and chromosomal fusion as a mechanism to heal broken chromosome ends. While this has been reported for cancer cells (Murnane 2012), it is generally considered a rare event in normal, non-transformed cells. Here we show that *de novo* telomere formation and chromosome fusion readily occur in a filamentous fungus during growth at temperature stress.

**Figure 3:**
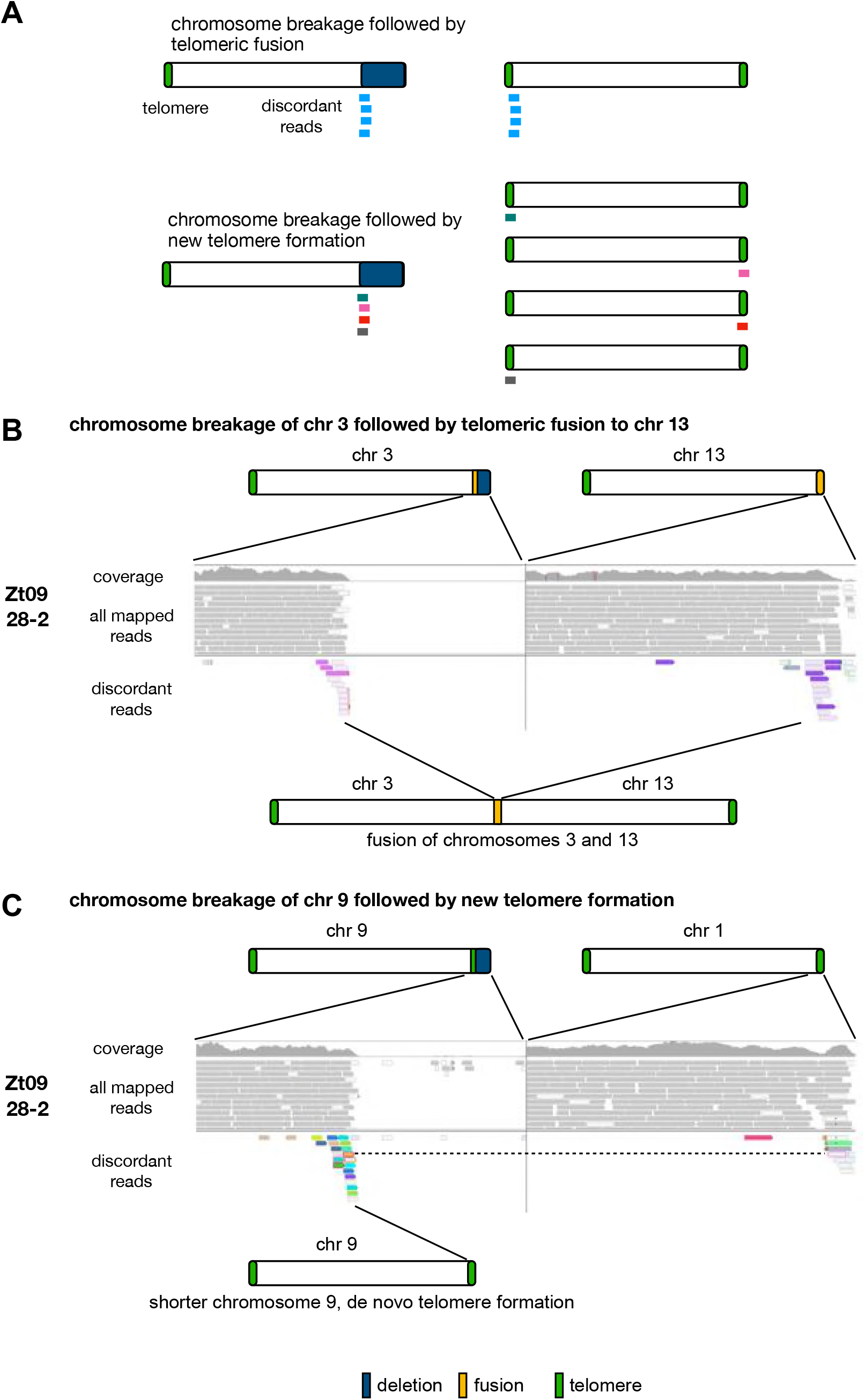
Discordant read analysis reveals chromosome fusion and chromosome breakage. A) Schematic illustration of discordant read mapping in the case of chromosome breakage and fusion. If chromosome breakage is followed by fusion to a different chromosome, the discordantly mapping reads at the breakpoint will have their respective read mate on the chromosome that is fused to the breakpoint. If breakage is followed by *de novo* telomere formation, the discordant reads have their respective read mates on telomeric repeats on random chromosomes. B and C) Read mapping to the genome of the reference isolate IPO323. B) In the experimentally evolved strains of *Z. tritici* at 28°C, the fusion of chromosomes 3 and 13 in the strain Zt09 28-2 is indicated by the increased occurrence of discordant reads at the chromosome breakpoint of the right arm of chromosome 3 and the right arm of chromosome 13. The color of the reads represents the chromosome their respective mate is mapping to. Color intensity of the reads indicates the mapping quality. C) New telomere formation at the breakpoint, here, as example, shown for chromosome 9 in the strain Zt09 28-2. Telomere formation is indicated by a high number of discordant reads, where the read mates are mapping to the telomeric repeats of different chromosomes. As an example, the right telomere of chromosome 1 is shown, where one mate of the discordant reads at the breakpoint of chromosome 9 is located (dashed line). A total of twelve breakage events that were followed by *de novo* telomere formation were detected in the five analyzed strains derived from the experiment at elevated temperature.

### Accessory chromosome instability also occurs in *Zymoseptoria* sister species

To assess whether the high frequency of mitotic chromosome loss is specific to *Z. tritici*, we conducted a short-term *in vitro* evolution experiment on the fungus *Z. ardabiliae*, a closely related sister species of *Z. tritici* that infects wild grasses. We used the previously characterized isolate STIR04 1.1.1 (Stukenbrock et al. 2011), here called Za17, in our experiments.

We first generated a high-quality reference genome based on long-read SMRT sequencing (Supplementary Text). To identify sequences of accessory chromosomes we first combined analyses of a population genomic dataset and PFGE (Stukenbrock et al. 2011; Stukenbrock & Dutheil 2018). Our analyses reveal at least four accessory chromosomes varying in their frequency among 17 *Z. ardabiliae* isolates. Based on these, we designed primers to amplify six loci located on putative accessory chromosomes of Za17 (Table S1).

Next, we conducted an *in vitro* evolution experiment at 18°C for four weeks in liquid culture with Za17 as the progenitor strain. Screening of 288 single clones by PCR identified five strains lacking one of the accessory chromosomes (^~^1.7%). We further verified the absence of accessory chromosomes by PFGE and whole genome sequencing for three of the five *Z. ardabiliae* strains (Figure 2B). These findings resemble the rapid loss of chromosomes in *Z. tritici* and suggest a common phenomenon, most likely related to mitotic cell division in the two *Zymoseptoria* species.

### Accessory chromosome losses affect *in planta* phenotypes

We next addressed the impact of spontaneous chromosome loss by comparing the fitness of the progenitor and evolved *Z. tritici* strains under different *in vitro* growth conditions, and during infection of a susceptible wheat variety by comparing growth and pycnidia formation. *In vitro*, the growth rate of strains lacking an additional accessory chromosome (Zt09A14 and Zt09A21) was comparable to the growth rate of the progenitor, Zt09, but with a slight tendency to slower growth of the derived chromosome-loss strains (Figure S2). For a more detailed phenotypic characterization, we used the five chromosome-loss strains isolated from pycnidia from the *in planta* experiment (Table S2). Several stress conditions including osmotic, oxidative, temperature, and cell wall stresses were tested *in vitro*. We compared the fungal phenotypes of the chromosome-loss strains and the progenitor strain IPO323 and observed no difference in growth rate or colony morphology between any chromosome-loss strain and IPO323 (Figure S3). *In planta*, however, we observed phenotypic differences between the evolved chromosome-loss strains and their progenitor. We measured fitness by counting the number of asexual fruiting bodies formed in stomata on infected wheat leaves and found a slightly higher fitness of the chromosome-loss strains compared to the progenitor strain (Figure 4). The difference was most pronounced for the strains lacking chromosome 14 and chromosome 19 where the density of pycnidia was found to be significantly higher (*p* values of 0.0003 and 0.015, Wilcoxon-Rank-Sum test and Holm’s i correction for multiple testing) compared to the IPO323 progenitor (Figure 4).

**Figure 4:**
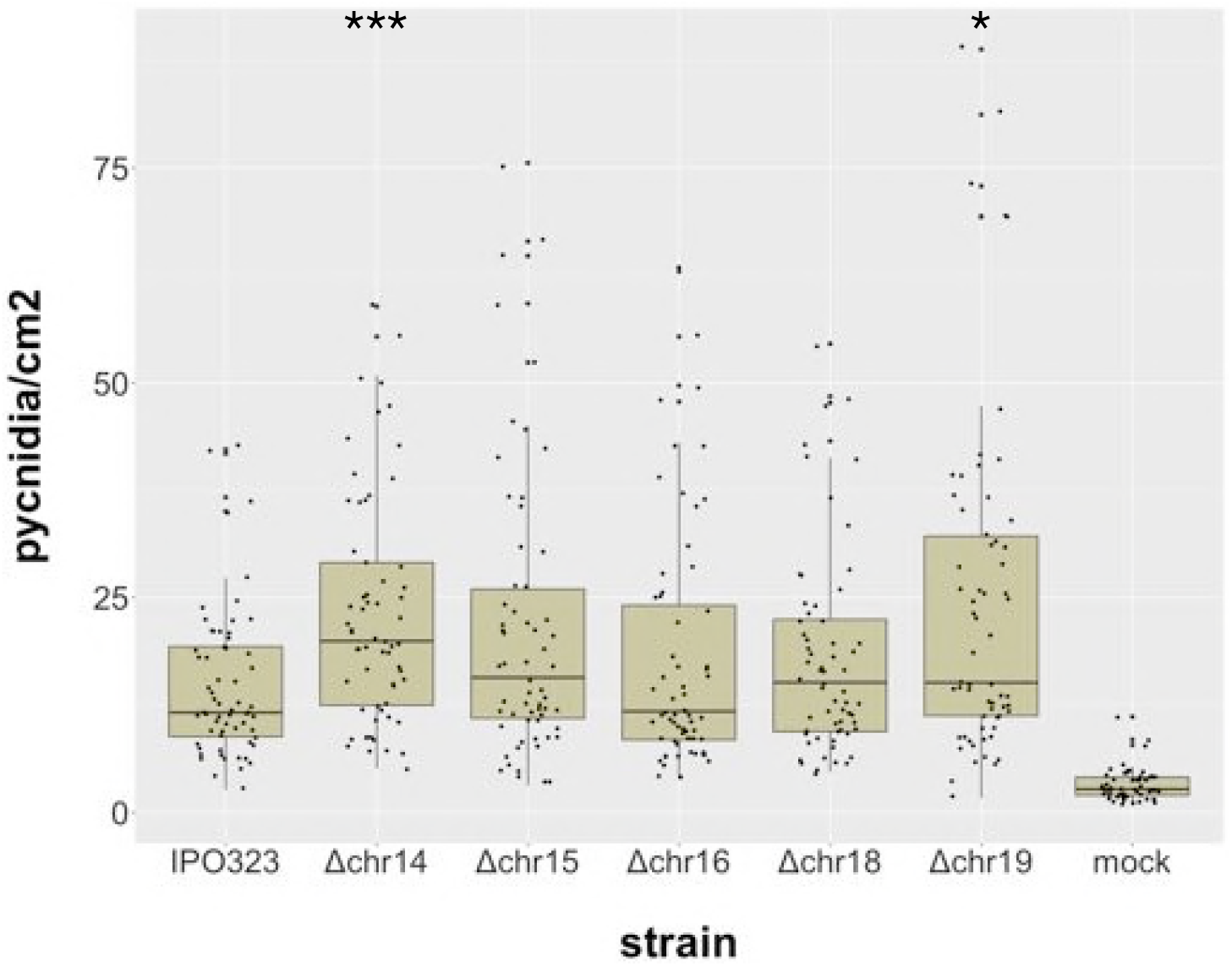
*In planta* infection assays reveal a negative effect of accessory chromosomes on fitness. To investigate the effect of accessory chromosome losses on *Z. tritici* infection of wheat, three independent experiments with the reference strain IPO323 and five chromosome-loss strains were conducted. The fitness of each strain was measured by counting the number of pycnidia per cm^2^ on the leaf surface. Statistical analysis (Wilcoxon-rank-sum test and Holms correction, p-values 0.0003 and 0.015) identified two chromosome-loss strains (Chr. 14 and 19) that show a significantly higher number of pycnidia compared to the reference strain IPO323 under the conditions used in this experiment.

## Discussion

### Consequences for evolution of accessory chromosomes

Accessory chromosomes were proposed to serve as an ‘evolutionary cradle’ for creating novel virulence genes without risking the disruption of essential genes on the core chromosomes (Croll & McDonald 2012). Indeed, signatures for accelerated evolution were shown by overall higher dN/dS ratios of genes on the accessory chromosomes compared to the core chromosomes (Stukenbrock et al. 2010). However, natural selection acts at the level of individuals and cannot maintain “a playground” for future beneficial effects. Rather, we hypothesize that the dynamic of accessory chromosome loss reflects a dynamic in the selective environment of *Z. tritici*, where under certain conditions these small chromosomes confer a fitness advantage, while under different conditions they cause decrease in fitness. A fitness advantage of chromosome-loss strains *in planta* may indicate the presence of avirulence factors on at least some accessory chromosomes. Avirulence factors can be recognized by the host to induce a resistance response resulting in reduction in virulence or even abortion of infection (Petit-Houdenot & Fudal 2017; Zhong et al. 2017). Our previous findings support this notion, as we found small but significant negative effects of accessory chromosomes during host infection (Habig et al. 2017). In other pathogenic fungi, however, the opposite effect has been demonstrated in several cases. A strong positive impact on virulence of accessory chromosomes have been observed in *Fusarium oxysporum* f. spp. and *Nectria haematococca* (Ma et al. 2010; Coleman et al. 2009; van Dam et al. 2018). In these pathogens chromosome losses resulted in complete loss of pathogenicity. While these cases are currently considered the norm, our observations show that responses to accessory chromosome loss can be more varied. Even though this fungus is an important pathogen in all wheat-growing countries, the biology of *Z. tritici* is still not completely understood. Mating and overwintering in the soil or litter layer remain largely unknown lifecycle stages, and we postulate that they impose different selection pressures on the fungus. During these stages, accessory chromosomes may confer significant fitness advantages that have not been demonstrated yet. Besides spontaneous chromosome loss during meiosis (Wittenberg et al. 2009), accessory chromosomes have been shown to follow non-Mendelian segregation and are transmitted at a significantly higher rate than expected, if one of the two mating partners lacks an accessory chromosome (Habig et al, submitted). While there is no comprehensive mechanistic explanation yet, chromosome conservation by re-replication during meiosis may counteract the chromosome loss during vegetative growth and therefore maintain accessory chromosomes in pathogen populations. In *Alternaria alternata*, the spontaneous loss of a conditionally dispensable chromosome during sub-culturing has been described for one isolate (Johnson et al. 2001). Furthermore, loss of a conditionally dispensable chromosome after meiosis has been observed in *Nectria haematococca* (aka *Fusarium solani* MPVI) (V. P. Miao et al. 1991). A recent study of the plant pathogenic fungus *Fusarium oxysporum* forma specialis *lycopersici* describes the spontaneous loss of dispensable chromosomes *in vitro* in one cell out of 35,000 (Vlaardingerbroek et al. 2016). This rate is much lower than the chromosome loss rate that we observe in *Z. tritici* and *Z. ardabiliae*, where, depending on the growth conditions, more than one cell out of two to one cell out of 50, lacks an accessory chromosome. In combination, these results indicate that chromosome instability is a common phenomenon in fungi, but the stability of chromosomes can vary substantially between different species. Studies in *Saccharomyces* and *Candida* species have shown that chromosome loss resulting in aneuploidy is a relatively frequent phenomenon in diploid cells and is advantageous under certain conditions (Kumaran et al. 2013; Bennett et al. 2014; Selmecki et al. 2010; Zhu et al. 2014). The loss of chromosomes in haploid organisms, such as *Z. tritici*, is expected to have more severe consequences, as genetic information is lost from the cell. We considered, however, that the effect of rapid chromosome loss or instability may be advantageous if these traits are selected for in pathogens like *Zymoseptoria*.

The temperature stress experiment *in vitro* showed that genome instability in *Z. tritici*, at this temperature, is not restricted to the accessory chromosomes, but also involves the core genome. New chromosome formation induced by telomere to telomere fusion followed by breakage of the dicentric chromosome was reported in *Cryptococcus neoformans* (Fraser et al. 2005) but likely occurred during meiosis rather than mitosis. In the asexual *Candida glabrata*, clinical isolates frequently inhabit chromosomal rearrangements indicating that structural variation acts as a virulence mechanism (Poláková et al. 2009) and might be a response to stresses such as antifungal treatment or elevated temperature as observed here. The instable regions close to the chromosome ends on the core chromosomes, however, show structural characteristics similar to the accessory chromosomes, such as higher repeat content, lower gene density and specific histone modification patterns (Dhillon et al. 2014; Grandaubert et al. 2015; Schotanus et al. 2015).

The accessory chromosomes of *Z. tritici* do not differ in terms of centromere or telomere organization from the core chromosomes, but the chromatin is largely transcriptionally inactive and potentially more condensed due to an enrichment with the histone H3 that is trimethylated at lysine 27 (H3K27me3), which covers almost all of the accessory chromosomes (Schotanus et al. 2015). H3K27me3 is also enriched in subtelomeric regions of core chromosomes, that we show here as prone to instability. High-resolution microscopic analyses indicate an unusual localization of centromeres in the *Z. tritici* nuclei, suggesting a distinct spatial organization of different chromosomes (Schotanus et al. 2015). Heterochromatin has been found close to the nuclear periphery, and H3K27me3 has been shown to be involved in facilitating lamina-proximal positioning (Harr et al. 2015). We hypothesize that the heterochromatic structure reflects a distinct physical organization of core and accessory chromosomes and likely the subtelomeric regions in the nucleus of *Z. tritici*, and that this is correlated to the instability of heterochromatic regions and chromosomes. A Hi-C study on *Neurospora crassa* showed that absence of H3K27me3 resulted in movement of subtelomeric regions and centromeres from the nuclear periphery into the nuclear matrix (Klocko et al. 2016). Further experiments focusing on localization of specific *Zymoseptoria* chromosomes in the nucleus by cytology or Hi-C (Galazka et al. 2016) should be conducted to address this hypothesis.

In conclusion, using experimental evolution, electrophoretic karyotyping and genome sequencing we showed that the accessory chromosomes of *Z. tritici* are highly unstable during asexual growth across numerous mitotic cell divisions (“mitotic growth”) *in vitro* as well as *in planta*. Surprisingly, increasing the temperature from 18 to 28°C dramatically increased overall genome instability in *Z. tritici*. Besides chromosome losses we observed structural variation in form of chromosome breakage, duplication, and fusion involving both core and accessory chromosomes, and all events were increased near telomeric sequences. In *Z. tritici*, all of these chromosome abnormalities have thus far been associated with meiosis (Croll et al. 2013; Wittenberg et al. 2009), however, our study highlights an important role of mitotic growth in generating genetic diversity. The variability in chromosome number and structure obtained over a relatively short period of time (during one infection event, or four weeks of vegetative growth) correlates with the genomic diversity observed in field populations of this important plant pathogen (Linde et al. 2002; Mehrabi et al. 2007; Zhan et al. 2003; McDonald & Martinez 1991). Our findings demonstrate that even sexual fungal pathogens can accelerate the generation of new genetic diversity by mitosis-associated structural variation. The mitotic events responsible for these rearrangements are still to be characterized.

Lastly, chromosome instability has been observed in various organisms and is a frequent phenomenon in cancer cells (McGranahan et al. 2012). Epigenetic factors and heterochromatin in particular have been shown to impact chromosome stability in human cancer cells (Slee et al. 2012). The accessory chromosome dynamics correlating with distinct chromatin patterns suggest that *Z. tritici* may provide a new model for mechanistic studies of chromosome instability in cancer cells.

## Materials and Methods

### Short term *in vitro* growth experiment in liquid culture

*Zymoseptoria* strains were diluted from glycerol stocks (−80°C), plated on YMS agar (4 g yeast extract, 4 g malt extract, 4 g sucrose, and 20 g agar per 1 liter) plates and grown for seven days at 18°C to obtain single colonies. One single colony was picked and suspended in 100 μL of YMS. Three replicate cultures were inoculated with 20 μL of cells from the single colony (^~^50,000 cells). Cells were grown in 25 mL YMS medium at 18°C or 28°C shaking at 200 rpm and 900 μL of the cultures were transferred to fresh medium after 3-4 days of growth. In total eight transfers were conducted, for a total time course of four weeks.

### Short term *in vitro* growth experiment on agar plates

A single colony derived directly from a plated dilution of frozen stock for Zt09 (IPO323A18) was resuspended in 1000 μL YMS including 25% glycerol by 2 min vortexing on a VXR basic Vibrax at 2,000 rpm, and 10-50 μL were replated onto a YMS agar plate. Forty replicates were produced. Cells were grown for seven days at 18°C whereby a random colony (based on vicinity to a prefixed position on the plate) derived from a single cell was picked and transferred to a new plate as described above. The transfer was conducted for a total of four times before a randomly chosen colony of each replicate was PCR screened and their complement on accessory chromosomes characterized as described below.

### Screening by PCR

Cells from transfer eight of liquid cultures were diluted and plated on YMS-agar plates to obtain single colonies. To extract DNA, single colonies were suspended in 50 μL of 25 mM NaOH and boiled at 98°C for 10 min; 50 μL of 40 mM Tris-HCl, pH 5.5, were added and 4 μL were used as template for the PCR. Primers for the right and left subtelomeric regions and close to the centromere were used for the chromosome loss screening in our *Z. tritici* isolates, for *Z. ardabiliae* primers in the center of candidate accessory chromosome unitigs were used. Primers and expected fragment lengths are listed in Table S1. Primers were designed with Clonemanager (Sci-Ed Software, Denver, USA) and MacVector and ordered from eurofins Genomics (Ebersberg, Germany).

### Plant experiments and pycnidia isolation

Growth conditions for plants were 16 h at light intensity of ^~^200 μmol/m^-2^s^-1^ and 8 h darkness in growth chambers at 20°C with 90% humidity. Seeds of the *Z. tritici* susceptible wheat cultivar Obelisk (Wiersum Plantbreeding BV, Winschoten, The Netherlands) were germinated on wet sterile Whatmann paper for four days at growth conditions before potting. Following potting, plants were further grown for seven days before infection.

For the phenotypic analyses, three independent experiments were conducted and 21 plants were used per strain per experiment. An ^~^5 cm long section on the second leaf of each plant was infected by brushing a cell suspension with 10^7^ cell/ml in H_2_O + 0.1% Tween 20 on the abaxial and adaxial side of the second leaf. Plants were placed into sealed bags containing ^~^1 L of water for 48 h to facilitate infection at maximum air humidity. Infected leaf areas were harvested 21 days post infection for further phenotypic analysis or prepared for pycnidia isolation by surface sterilization with 1.2% NaClO for 2 min followed by 70% ethanol for a few seconds and washed twice with H_2_O. Leaves were placed into a sterile environment with maximum air humidity and incubated for 7-14 days at plant growth conditions. Spores that had been pressed out from pycnidia were isolated using a sterile syringe needle and resuspended into 50 μL YMS medium with 50% glycerol. Cells were suspended by 30 min vortexing on a VXR basic Vibrax (IKA, Staufen, Germany) at 2,000 rpm, plated on YMS agar, and incubated for seven days at 18°C. Colonies arising from single cells were PCR screened and their complement on accessory chromosomes characterized as described below.

### Phenotypic characterization of infected leaves

Harvested leaves were taped to a sheet of paper and scanned using a flatbed scanner at a resolution of 2,400 dpi. The scanned leaves were analyzed using ImageJ (Schneider et al. 2012) and a plug-in described previously (Stewart et al. 2016). The measurement of pycnidia/cm^2^ was used for further statistical analyses in R (Ihaka & Gentleman 1996). The Wilcoxon-Rank-Sum test and the Holm’s correction for multiple testing were applied to assess statistical differences between the samples.

### *In vitro* phenotype assay

A spore solution containing 10^7^ cells/mL and tenfold dilution series to 1,000 cells/mL was prepared. To test for responses to different stress conditions *in vitro*, YMS plates containing NaCl (0.5 M and 1 M), sorbitol (1 M and 1.5 M), Congo Red (300 μg/mL and 500 μg/ml), H_2_O_2_ (1.5 mM and 2 mM), MMS (methyl methanesulfonate, 0.01 %), a H_2_O-agar plate and two plates containing only YMS were prepared. Three μL of the spore suspension dilutions were pipetted on the plates and incubated at 18°C for seven days. One of the YMS plates was incubated at 28°C to test for thermal stress responses.

### *In vitro* growth assay

YMS liquid cultures containing 10^5^ cells/mL of either Zt09 or the chromosome-loss strains Zt09Δ14 and Zt09Δ21 were prepared. Three replicates per strain were used in each experiment. The spores were grown for four days at 18°C at 200 rpm in 25 mL YMS and the OD_600_ was measured at different time points throughout the experiment. The data was fitted using the R package growthcurver (Sprouffske & Wagner 2016) and the r values of each replicate used for statistical comparison. Wilcoxon rank-sum test was used to compare the r values of the different strains.

### Pulsed field gel electrophoresis (PFGE) and Southern blotting

Fungal strains were grown in YMS medium for five days. Cells were harvested by centrifugation for 10 min at 3,500 rpm. 5 × 10^8^ cells were used for plug preparation. The cells were resuspended in 1 mL H_2_O and mixed with 1 mL of 2.2 % low range ultra agarose (BioRad, Munich, Germany). The mixture was pipetted into plug casting molds and cooled for 1 h at 4°C. The plugs were transferred to 50 mL screw cap Falcon tubes containing 5 mL of lysis buffer (1 % SDS; 0.45 M EDTA; 1.5 mg/ml proteinase K, Roth, Karlsruhe, Germany), and incubated in lysis buffer for 48 h at 55°C, replacing the buffer once after 24 h. Chromosomal plugs were washed three times for 20 min with 1 × TE buffer before storage in 5 mL of 0.5 M EDTA at 4°C. PFGE was performed with a CHEF-DR III pulsed field electrophoresis system (BioRad, Munich, Germany). To separate the small accessory chromosomes, the following settings were applied: switching time 50 s – 150 s, 5 V/cm, 120° angle, 1 % pulsed field agarose (BioRad, Munich, Germany) in 0.5 × TBE (TRIS/borate/EDTA) for 48 h. Separation of mid-size chromosomes was conducted with the settings: switching time 250 s – 1000 s, 3 V/cm, 106° angle, 1 % pulsed field agarose in 0.5 × TBE for 72 h. *Saccharomyces cerevisiae* chromosomal DNA (BioRad, Munich, Germany) was used as size marker for the accessory chromosomes, and *Hansenula wingei* chromosomal DNA (BioRad, Munich, Germany) for mid-size chromosomes. Gels were stained in ethidium bromide staining solution (1 μg/ml ethidium bromide in H_2_O) for 30 min. Detection of chromosomal bands was performed with the GelDocTM XR+ system (Bio-Rad, Munich, Germany).

Southern blotting was performed as described previously (Southern 1975) using DIG-labeled probes generated with the PCR DIG labeling Mix (Roche, Mannheim, Germany) following the manufacturer’s instructions.

### Sequencing and genome comparison

DNA for whole genome sequencing was prepared as described in (Allen et al. 2006). Library preparation with a reduced number of PCR cycles (four cycles) and sequencing were performed by the Max Planck Genome Center, Cologne, Germany (http://mpgc.mpipz.mpg.de/home/) using an Illumina HiSeq3000 machine, obtaining ^~^30x coverage with 150-nt paired-end reads. Raw reads were quality filtered using Trimmomatic and mapped to the reference genome of *Z. tritici* or *Z. ardabiliae* using bowtie2. SNP calling was conducted with samtools and bcftools and further quality filtered. The filtered SNPs were validated by manual inspection. The data was visualized in IGV (Integrative Genomics Viewer, http://software.broadinstitute.org/software/igv/) (Thorvaldsdóttir et al. 2013). Detailed commands for each step of the data processing pipeline can be found in the Supplementary Text.

### Accession numbers

The sequencing data generated for this project were deposited in the Sequence Read Archive under BioProject ID PRJNA428438.

## Acknowledgements

Marcello Zala, Daniel Croll and Christoph J. Eschenbrenner are acknowledged for the preparation and assembly of the SMRT sequencing. Kathrin Happ for assistance during plant experiments. We thank all current and past members of the Environmental Genomics Group for fruitful discussions and overall support. Research in the group of EHS is supported by the Max-Planck Society, the state of Schleswig-Holstein and the DFG priority program SPP1819.

## Author contributions

MM: Conceptualization, Investigation, Visualization, Formal analysis, Writing-original draft

MH: Conceptualization, Investigation (*in planta* chromosome loss), Writing—review & editing

MF: Conceptualization, Formal analysis, Writing—review & editing, Supervision

EHS: Conceptualization, Formal analysis, Writing—review & editing, Supervision, Project administration, Funding acquisition

## Supplement

**Table S1:**
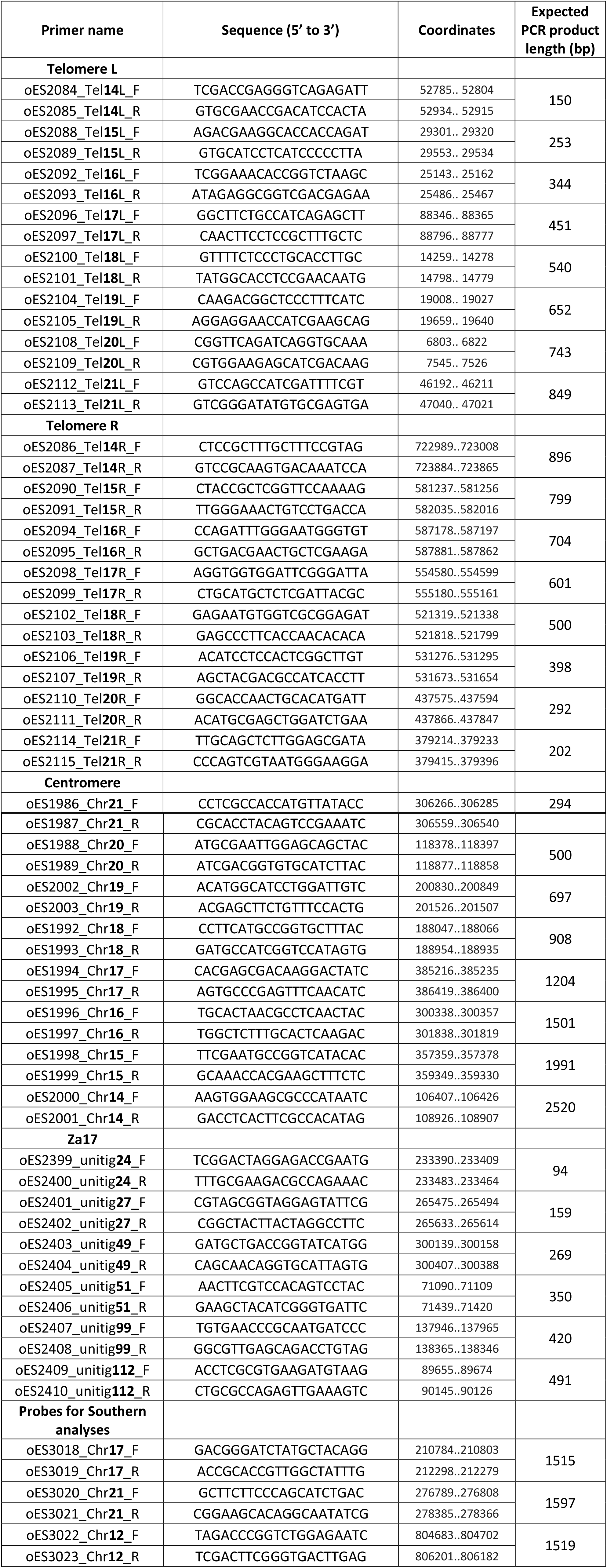
List of primers used for screening for accessory chromosomes in *Z. tritici* and *Z. ardabiliae*.

**Table S2:** List of chromosome-loss strains identified in the *in vitro* and *in planta* experiments. Strains described in detail in the main text are highlighted in **bold**.

| Experiment | Strain name | Chromosome(s) lost | Strain name in main text | Method used to confirm chromosome loss |
| --- | --- | --- | --- | --- |
| progenitor | **Zt09** | **18** | **Zt09<sup>3)</sup>** | **PCR, PFGE<sup>1)</sup>, WGS<sup>2)</sup>** |
| progenitor | **IPO323** | - | **IPO323<sup>4)</sup>** | **PCR, PFGE, WGS** |
| progenitor | **STIR04 1.1.1** | - | **Za17** | **PCR, PFGE, WGS** |
| **Zt09 *in vitro* liquid culture 18°C** |  |  |  |  |
|  | **Zt09_18_1.1.38** | **14 (18)** | **Zt09Δ14<sup>3)</sup>** | **PCR, PFGE, WGS** |
|  | **Zt09_18_1.1.45** | **15 (18)** | **Zt09Δ15** | **PCR, PFGE, WGS** |
|  | Zt09_18_1.1.51 | 14 (18) |  |  |
|  | **Zt09_18_1.1.71** | **21 (18)** | **Zt09Δ21<sup>3)</sup>** | **PCR, PFGE, WGS** |
|  | Zt09_18_1.1.85 | 16 (18) |  | PCR, PFGE |
|  | Zt09_18_1.1.95 | 14 (18) |  | PCR, PFGE |
|  | Zt09_18_2.1.3 | 16 (18) |  | PCR, PFGE |
|  | Zt09_18_2.1.4 | 14 (18) |  | PCR, PFGE |
|  | Zt09_18_2.1.21 | 15 (18) |  | PCR, PFGE |
|  | Zt09_18_2.1.24 | 15 (18) |  | PCR, PFGE |
|  | Zt09_18_2.1.33 | 14 (18) |  | PCR, PFGE |
|  | Zt09_18_2.1.64 | 14 (18) |  | PCR, PFGE |
|  | Zt09_18_2.1.68 | 16 (18) |  | PCR, PFGE |
|  | Zt09_18_2.1.73 | 20 (18) |  | PCR, PFGE |
|  | Zt09_18_2.1.79 | 16 (18) |  | PCR, PFGE |
|  | Zt09_18_2.1.85 | 14 (18) |  | PCR, PFGE |
|  | Zt09_18_1.2.4 | 15 (18) |  | PCR, PFGE |
|  | **Zt09_18_1.2.18** | **20 (18)** | **Zt09Δ20** | **PCR, PFGE, WGS** |
|  | Zt09_18_1.2.40 | 15 (18) |  | PCR |
|  | Zt09_18_1.2.48 | 15 (18) |  | PCR |
|  | Zt09_18_1.2.86 | 15 (18) |  | PCR |
|  | Zt09_18_1.2.89 | 16 (18) |  | PCR |
|  | Zt09_18_1.2.93 | 14 (18) |  | PCR |
|  | Zt09_18_2.2.6 | 14 (18) |  | PCR |
|  | Zt09_18_2.2.38 | 14 (18) |  | PCR |
|  | Zt09_18_2.2.43 | 15 (18) |  | PCR |
|  | Zt09_18_2.2.64 | 14 (18) |  | PCR |
|  | Zt09_18_2.2.70 | 16 (18) |  | PCR |
|  | Zt09_18_2.2.74 | 16 (18) |  | PCR |
|  | Zt09_18_1.3.40 | 14 (18) |  | PCR |
|  | Zt09_18_1.3.41 | 14 (18) |  | PCR |
|  | Zt09_18_1.3.92 | 16 (18) |  | PCR |
|  | Zt09_18_2.3.21 | 14 (18) |  | PCR |
|  | Zt09_18_2.3.41 | 14 (18) |  | PCR |
|  | Zt09_18_2.3.49 | 14 (18) |  | PCR |
|  | Zt09_18_2.3.53 | 14 (18) |  | PCR |
|  | Zt09_18_2.3.85 | 14 (18) |  | PCR |
|  | Zt09_18_2.3.86 | 16 (18) |  | PCR |
| **Zt09 *in vitro***<br>**liquid culture**<br>**28°C** |  |  |  |  |
|  | Zt09_28_1.1.1 | 14 (18) |  | PCR |
|  | Zt09_28_1.1.2 | 14 (18) |  | PCR |
|  | Zt09_28_1.1.4 | 14 (18) |  | PCR |
|  | Zt09_28_1.1.5 | 14 (18) |  | PCR |
|  | Zt09_28_1.1.6 | 14 (18) |  | PCR |
|  | Zt09_28_1.1.8 | 14 (18) |  | PCR |
|  | Zt09_28_1.1.10 | 14 (18) |  | PCR |
|  | Zt09_28_1.1.11 | 14,21 (18) |  | PCR, PFGE |
|  | Zt09_28_1.1.12 | 14,20 (18) |  | PCR, PFGE |
|  | Zt09_28_1.1.13 | 14 (18) |  | PCR |
|  | Zt09_28_1.1.14 | 14 (18) |  | PCR |
|  | Zt09_28_1.1.15 | 14 (18) |  | PCR |
|  | Zt09_28_1.1.17 | 14 (18) |  | PCR |
|  | Zt09_28_1.1.19 | 14,20,21 (18) | **Zt09 28-1** | **PCR, PFGE, WGS** |
|  | Zt09_28_1.1.21 | 14 (18) |  | PCR |
|  | Zt09_28_2.1.2 | 19 (18) |  | PCR |
|  | Zt09_28_2.1.3 | 14 (18) |  | PCR |
|  | Zt09_28_2.1.4 | 14 (18) |  | PCR |
|  | Zt09_28_2.1.6 | 14 (18) |  | PCR |
|  | Zt09_28_2.1.7 | 14 (18) |  | PCR |
|  | Zt09_28_2.1.8 | 14 (18) |  | PCR |
|  | Zt09_28_2.1.9 | 14 (18) |  | PCR |
|  | Zt09_28_2.1.10 | 14 (18) |  | PCR |
|  | Zt09_28_2.1.11 | 14 (18) |  | PCR |
|  | Zt09_28_2.1.12 | 14 (18) |  | PCR |
|  | Zt09_28_2.1.14 | 14,20 (18) |  | PCR |
|  | Zt09_28_2.1.15 | 14 (18) |  | PCR |
|  | Zt09_28_2.1.16 | 14 (18) |  | PCR |
|  | Zt09_28_2.1.17 | 14 (18) |  | PCR |
|  | Zt09_28_2.1.18 | 14 (18) |  | PCR |
|  | Zt09_28_2.1.19 | 14 (18) |  | PCR |
|  | Zt09_28_2.1.21 | 14,20,21 (18) |  | PCR |
|  | Zt09_28_2.1.22 | 20 (18) |  | PCR |
|  | Zt09_28_2.1.24 | 14,16 (18) |  | PCR |
|  | Zt09_28_2.1.25 | 14 (18) |  | PCR |
|  | Zt09_28_2.1.26 | 14,20 (18) |  | PCR |
|  | Zt09_28_2.1.27 | 14 (18) |  | PCR |
|  | Zt09_28_2.1.28 | 14 (18) |  | PCR |
|  | Zt09_28_2.1.29 | 14 (18) |  | PCR |
|  | Zt09_28_2.1.31 | 14 (18) |  | PCR |
|  | Zt09_28_2.1.33 | 14 (18) |  | PCR |
|  | Zt09_28_2.1.34 | 14 (18) |  | PCR |
|  | Zt09_28_2.1.35 | 14 (18) |  | PCR |
|  | Zt09_28_2.1.37 | 14 (18) |  | PCR |
|  | Zt09_28_2.1.38 | 14,20 (18) |  | PCR, PFGE |
|  | Zt09_28_2.1.39 | 21 (18) |  | PCR, PFGE |
|  | Zt09_28_2.1.40 | 14 (18) |  | PCR |
|  | Zt09_28_2.1.41 | 14 (18) |  | PCR |
|  | Zt09_28_2.1.42 | 14 (18) |  | PCR |
|  | Zt09_28_1.2.1 | 14,20 (18) |  | PCR |
|  | Zt09_28_1.2.2 | 14 (18) |  | PCR |
|  | Zt09_28_1.2.3 | 14 (18) |  | PCR |
|  | Zt09_28_1.2.4 | 14,15,16,19,20 (18) | **Zt09 28-2** | **PCR, PFGE, WGS** |
|  | Zt09_28_1.2.5 | 14 (18) |  | PCR |
|  | Zt09_28_1.2.6 | 14 (18) |  | PCR |
|  | Zt09_28_1.2.7 | 14,16 (18) | **Zt09 28-3** | **PCR, PFGE, WGS** |
|  | Zt09_28_1.2.8 | 14 (18) |  | PCR |
|  | Zt09_28_1.2.9 | 14 (18) |  | PCR |
|  | Zt09_28_1.2.10 | 14 (18) |  | PCR |
|  | Zt09_28_1.2.11 | 14 (18) |  | PCR |
|  | Zt09_28_1.2.12 | 14,15 (18) |  | PCR, PFGE |
|  | Zt09_28_1.2.15 | 14 (18) |  | PCR |
|  | Zt09_28_1.2.16 | 14 (18) |  | PCR |
|  | Zt09_28_1.2.17 | 14 (18) |  | PCR |
|  | Zt09_28_1.2.18 | 14,15,20,21 (18) | **Zt09 28-4** | **PCR, PFGE, WGS** |
|  | Zt09_28_1.2.19 | 14,16 (18) |  | PCR, PFGE |
|  | Zt09_28_1.2.20 | 14,21 (18) |  | PCR, PFGE |
|  | Zt09_28_1.2.21 | 14 (18) |  | PCR |
|  | Zt09_28_1.2.22 | 14 (18) |  | PCR |
|  | Zt09_28_1.2.23 | 14,20 (18) |  | PCR, PFGE |
|  | Zt09_28_1.2.24 | 14 (18) |  | PCR |
|  | Zt09_28_1.2.25 | 14 (18) |  | PCR |
|  | Zt09_28_1.2.26 | 14 (18) |  | PCR |
|  | Zt09_28_1.2.27 | 14,15 (18) |  | PCR, PFGE |
|  | Zt09_28_1.2.28 | 14 (18) |  | PCR |
|  | Zt09_28_1.2.29 | 14 (18) |  | PCR |
|  | Zt09_28_1.2.30 | 14 (18) |  | PCR |
|  | Zt09_28_1.2.31 | 14 (18) |  | PCR |
|  | Zt09_28_1.2.32 | 14 (18) |  | PCR |
|  | Zt09_28_2.2.1 | 14 (18) |  | PCR |
|  | Zt09_28_2.2.2 | 14 (18) |  | PCR |
|  | Zt09_28_2.2.3 | 15 (18) |  | PCR |
|  | Zt09_28_2.2.4 | 14 (18) |  | PCR |
|  | Zt09_28_2.2.5 | 14 (18) |  | PCR |
|  | Zt09_28_2.2.6 | 15 (18) |  | PCR |
|  | Zt09_28_2.2.7 | 14 (18) |  | PCR |
|  | Zt09_28_2.2.10 | 14 (18) |  | PCR |
|  | Zt09_28_2.2.11 | 14 (18) |  | PCR |
|  | Zt09_28_2.2.12 | 14,16 (18) |  | PCR |
|  | Zt09_28_2.2.13 | 14 (18) |  | PCR |
|  | Zt09_28_2.2.14 | 14,16 (18) |  | PCR |
|  | Zt09_28_2.2.15 | 14,20 (18) |  | PCR |
|  | Zt09_28_2.2.16 | 14,15 (18) | **Zt09 28-5** | **PCR, PFGE, WGS** |
|  | Zt09_28_2.2.17 | 14 (18) |  | PCR |
|  | Zt09_28_2.2.18 | 14,16 (18) |  | PCR |
|  | Zt09_28_2.2.19 | 14 (18) |  | PCR |
|  | Zt09_28_2.2.20 | 14 (18) |  | PCR |
|  | Zt09_28_2.2.21 | 14 (18) |  | PCR |
|  | Zt09_28_2.2.22 | 14,15 (18) |  | PCR |
|  | Zt09_28_2.2.23 | 14 (18) |  | PCR |
|  | Zt09_28_2.2.24 | 14 (18) |  | PCR |
|  | Zt09_28_2.2.25 | 14 (18) |  | PCR |
|  | Zt09_28_2.2.26 | 14,20 (18) |  | PCR |
|  | Zt09_28_2.2.27 | 19 (18) |  | PCR |
|  | Zt09_28_2.2.28 | 14 (18) |  | PCR |
|  | Zt09_28_1.3.1 | 14 (18) |  | PCR |
|  | Zt09_28_1.3.2 | 14 (18) |  | PCR |
|  | Zt09_28_1.3.4 | 14,20 (18) |  | PCR, PFGE |
|  | Zt09_28_1.3.5 | 16,20 (18) |  | PCR, PFGE |
|  | Zt09_28_1.3.6 | 14 (18) |  | PCR |
|  | Zt09_28_1.3.7 | 14,16 (18) |  | PCR, PFGE |
|  | Zt09_28_1.3.8 | 14,20 (18) |  | PCR |
|  | Zt09_28_1.3.11 | 14 (18) |  | PCR |
|  | Zt09_28_1.3.13 | 14 (18) |  | PCR |
|  | Zt09_28_1.3.14 | 14,20,21 (18) |  | PCR |
|  | Zt09_28_1.3.15 | 14 (18) |  | PCR |
|  | Zt09_28_1.3.17 | 14 (18) |  | PCR |
|  | Zt09_28_1.3.18 | 14 (18) |  | PCR |
|  | Zt09_28_1.3.19 | 14 (18) |  | PCR |
|  | Zt09_28_1.3.20 | 14 (18) |  | PCR |
|  | Zt09_28_1.3.22 | 14 (18) |  | PCR |
|  | Zt09_28_1.3.24 | 14,15 (18) |  | PCR |
|  | Zt09_28_1.3.27 | 14 (18) |  | PCR |
|  | Zt09_28_1.3.28 | 14 (18) |  | PCR |
|  | Zt09_28_2.3.1 | 14,20 (18) |  | PCR |
|  | Zt09_28_1.3.3 | 14 (18) |  | PCR |
|  | Zt09_28_1.3.5 | 14 (18) |  | PCR |
|  | Zt09_28_1.3.6 | 14 (18) |  | PCR |
|  | Zt09_28_1.3.7 | 14 (18) |  | PCR |
|  | Zt09_28_1.3.9 | 14 (18) |  | PCR |
|  | Zt09_28_1.3.11 | 14 (18) |  | PCR |
|  | Zt09_28_1.3.12 | 14 (18) |  | PCR |
|  | Zt09_28_1.3.13 | 14 (18) |  | PCR |
|  | Zt09_28_1.3.14 | 14,21 (18) |  | PCR |
|  | Zt09_28_1.3.15 | 14 (18) |  | PCR |
|  | Zt09_28_1.3.17 | 14 (18) |  | PCR |
|  | Zt09_28_1.3.19 | 14 (18) |  | PCR |
|  | Zt09_28_1.3.20 | 14 (18) |  | PCR |
|  | Zt09_28_1.3.22 | 14 (18) |  | PCR |
|  | Zt09_28_1.3.26 | 14 (18) |  | PCR |
|  | Zt09_28_1.3.27 | 14,15 (18) |  | PCR |
|  | Zt09_28_1.3.28 | 14 (18) |  | PCR |
|  | Zt09_28_1.3.29 | 14 (18) |  | PCR |
|  | Zt09_28_1.3.30 | 14 (18) |  | PCR |
|  | Zt09_28_1.3.32 | 14 (18) |  | PCR |
|  | Zt09_28_1.3.33 | 14,19 (18) |  | PCR |
|  | Zt09_28_1.3.35 | 14 (18) |  | PCR |
|  | Zt09_28_1.3.36 | 14 (18) |  | PCR |
| **Zt09 *in vitro***<br>**on plate 18°C** |  |  |  |  |
|  | 2-08-005 | 16 (18) |  | PCR |
|  | 2-09-005 | 16 (18) |  | PCR |
|  | 2-14-005 | 16 (18) |  | PCR |
|  | 2-17-005 | 16 (18) |  | PCR |
|  | 2-01-005 | 21 (18) |  | PCR |
| **IPO323 *in planta*** |  |  |  |  |
|  | **A#101** | **14** | **IPO323Δ14<sup>4)</sup>** | **PCR, PFGE, WGS** |
|  | A#52 | 15 |  | PCR |
|  | **B#56** | **15** | **IPO323Δ15<sup>4)</sup>** | **PCR, PFGE, WGS** |
|  | B#181 | 15 |  | PCR |
|  | PycB F15 13 | 15 |  | PCR |
|  | PycB F15 52 | 15 |  | PCR |
|  | PycB F30 6 | 15 |  | PCR |
|  | **PycB E 29 57** | **16** | **IPO323Δ16<sup>4)</sup>** | **PCR** |
|  | PycA E10#17 | 18 |  | PCR |
|  | **B#152** | **18** | **IPO323Δ18<sup>4)</sup>** | **PCR, PFGE, WGS** |
|  | PycB F15 1 | 18 |  | PCR |
|  | PycB F15 75 | 18 |  | PCR |
|  | PycB F15 83 | 18 |  | PCR |
|  | PycB F30 43 | 18 |  | PCR |
|  | PycB F30 90 | 18 |  | PCR |
|  | **B#59** | **19** | **IPO323Δ19<sup>4)</sup>** | **PCR, PFGE, WGS** |
|  | PycB F15 79 | 19 |  | PCR |
| **Za17 *in vitro***<br>**liquid culture**<br>**18°C** |  |  |  |  |
|  | **Za17_2.46** | **unitig 24** | **ZaΔ24** | **PCR, PFGE, WGS** |
|  | **Za17_2.61** | **unitig 49** | **ZaΔ49** | **PCR, PFGE, WGS** |
|  | **Za17_2.69** | **unitig 51** | **ZaΔ51** | **PCR, PFGE, WGS** |
|  | Za17_2.81 | unitig 49 |  | PCR, PFGE |
|  | Za17_3.38 | unitig 51 |  | PCR, PFGE |

<sup>1)</sup> Pulsed-field gel electrophoresis
<sup>2)</sup> Whole genome sequencing
<sup>3)</sup> Strains used for *in vitro* growth assay
<sup>4)</sup> Strains used for *in vitro* and *in planta* phenotyping

**Table S3:** Chromosome variation of sequenced Zt09-derived strains grown at 28°C. Absent sequences at the ends of each chromosome (left (L) or right (R) arm) are indicated, as well as the complete absence or a du

| Chromosome | Zt09 28-1 | Zt09 28-2 | Zt09 28-3 | Zt09 28-4 | Zt09 28-5 |
| --- | --- | --- | --- | --- | --- |
| 1 |  |  | <sup>2)</sup> internal tel<br>183,500 |  |  |
| 2 |  |  | L (5 kb) new tel 2<br>breakpoints<br>R (4 kb) new tel |  |  |
| 3 |  | R (10 kb)<br>attached to chr 13 |  |  |  |
| 4 |  |  |  |  | L (10 kb)<br>new tel |
| 5 | R (35 kb)<sup>1)</sup><br>new tel | R (35 kb)<sup>1)</sup><br>new tel | R (35 kb)<sup>1)</sup><br>new tel | R (35 kb)<sup>1)</sup><br>new tel | R (35 kb)<sup>1)</sup><br>new tel |
| 6 |  |  | R (2 kb) new tel |  |  |
| 7 |  |  |  |  |  |
| 8 | R (200 bp) |  |  |  |  |
| 9 | R (16 kb)<br>new tel | R (3 kb)<br>new tel |  |  |  |
| 10 |  |  | L (27 kb) new tel |  | L (7 kb) |
| 11 |  |  | R (2.5 kb) new tel |  |  |
| 12 |  |  |  |  |  |
| 13 |  | R<br>attached to chr 3 |  | R (1.5 kb)<br>new tel |  |
| 14 | absent | absent | absent | absent | absent |
| 15 | R (350 bp)<br>new tel | absent |  | absent | absent |
| 16 |  | absent | absent |  |  |
| 17 |  | **duplicated** | L (60 kb) new tel<br>R (1kb) | **duplicated**<br><sup>2)</sup> internal<br>tel 54,000 |  |
| 18 | absent | absent | absent | absent | absent |
| 19 |  | absent | L (22 kb) new tel |  |  |
| 20 | absent | absent |  | absent | **duplicated** |
| 21 | absent | L (400 bp)<br><sup>2)</sup> internal tel<br>32,000 |  | absent |  |

<sup>1)</sup> The 35 kb sequence missing on the right arm of chromosome five is already absent in the progenitor strain Zt09. Discordant reads suggest a new telomere formation indicated as 'new tel'.
<sup>2)</sup> Coordinates of cases where new telomeres were not located at chromosome breakpoints.

**Table S4:**
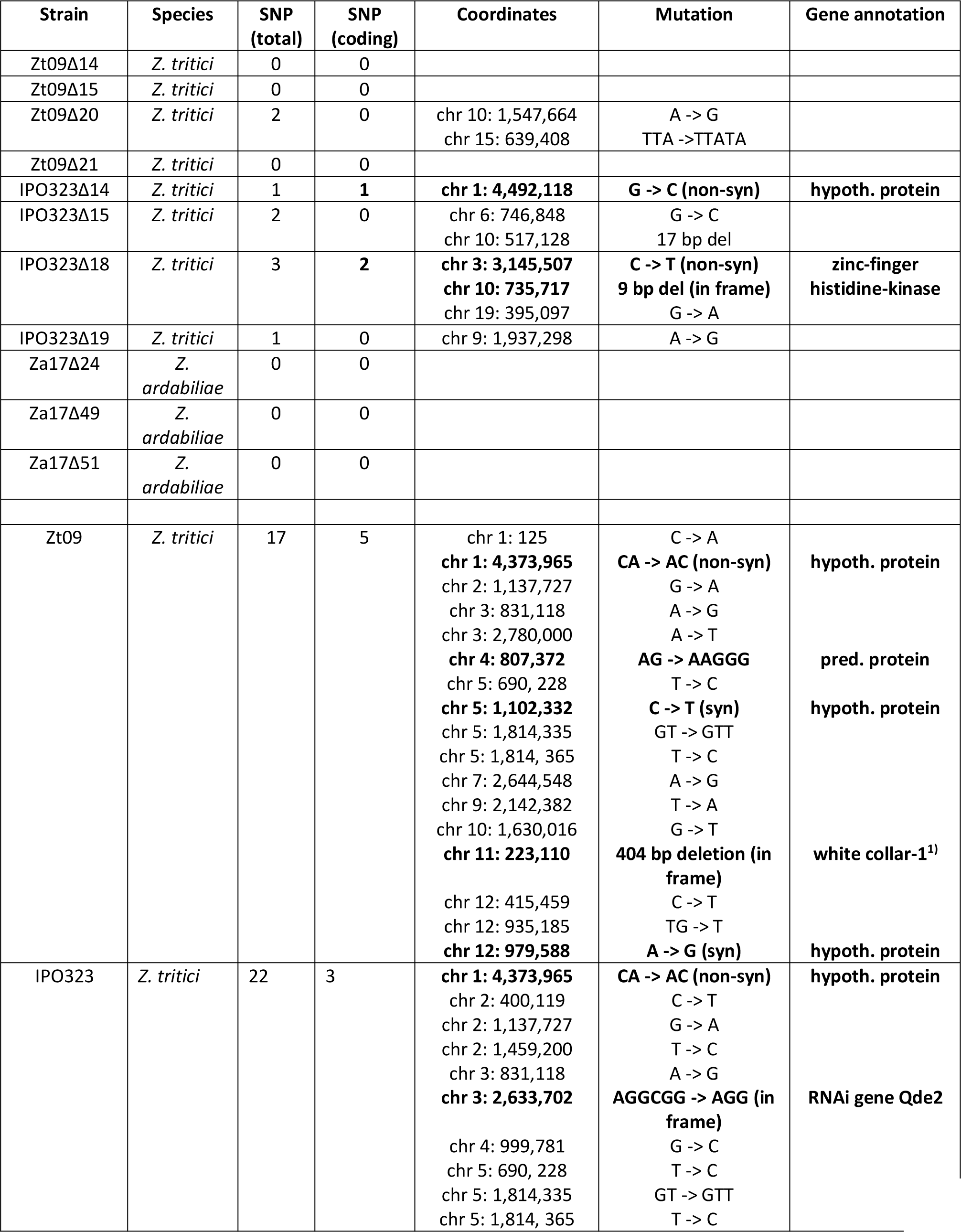
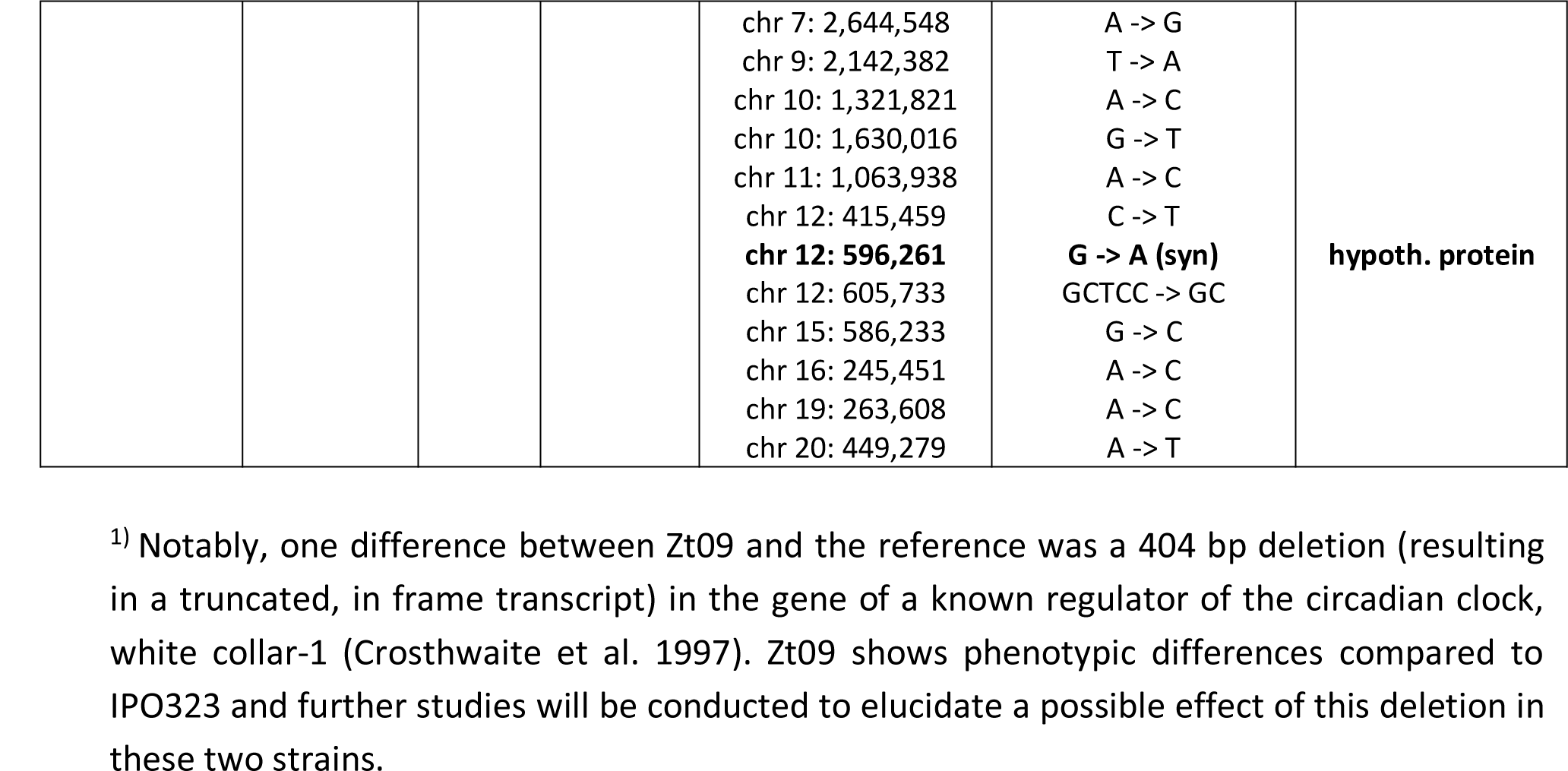
Location and annotation of SNPs and INDELs found in the sequenced chromosome-loss strains. Listed are SNPs that deviate in the chromosome loss strains from the respective corresponding ancestral strain Zt09, IPO323 or Za17. Also listed are the SNPs that differ between the ancestral strains Zt09 and IPO323 and the reference genome. Polymorphisms in coding regions are indicated in **bold**.

**Figure S1:**
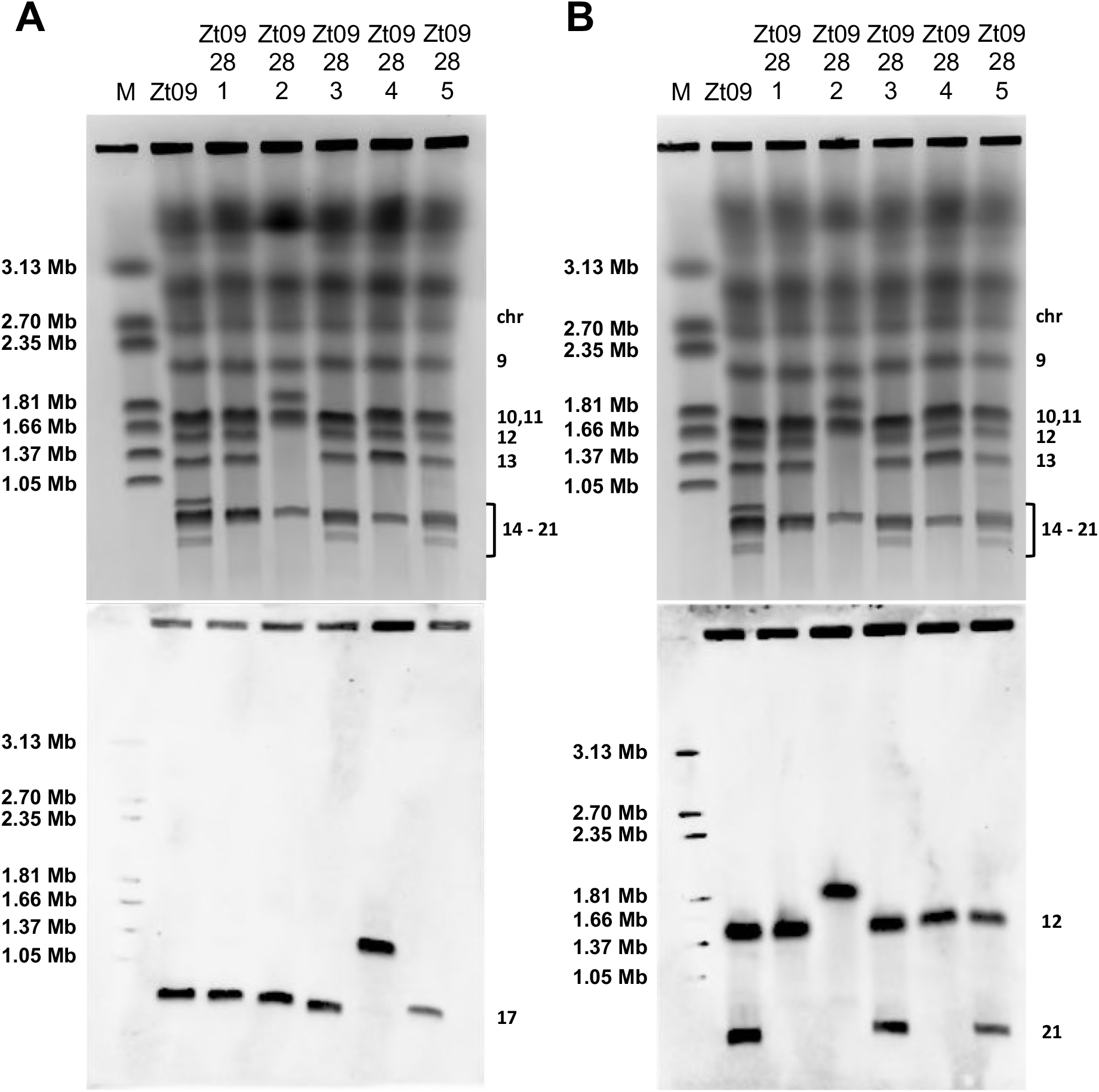
Southern blotting of pulsed-field gels using chromosome-specific probes confirms chromosome fusions. To prove and visualize chromosome fusions in the strains derived from the temperature stress experiment, we conducted PFGE followed by Southern blots with probes specific to the chromosomes, which based on genome data and karyotype analyses, were found to have fused. Panel (A) shows a color-inverted image of the pulsed-field gel (top) and Southern blot (bottom) using a probe specific for chromosome 17. The genomes of all tested strains have chromosome 17, and the genomes of strains Zt09 28-2 and Zt09 28-4 contain a duplicated chromosome 17 (Figure 2C). While in Zt09 28-2 the duplication resulted in two separate copies of the chromosome, the two chromosomes 17 in Zt09 28-4 fused and formed a new ^~^ 1.2 Mb chromosome. (B) For the second Southern blot, we used probes for chromosomes 12 and 21. Chromosome 21 is absent in the strains Zt09 28-1 and Zt09 28-3 (PFGE, top and Southern analyses, bottom). Genome sequencing shows that chromosome 21 is present in Zt09 28-2 (Figure 2C), however the respective band on the pulsed-field gel is missing. Similarly, chromosomes 12 and 13 are not present in this strain (PFGE, top), while the chromosome sequences are present in the whole genome sequence data. Apparent loss of chromosome 13 can be explained by the fusion of chromosomes 3 and 13, as indicated by the sequence analysis (Figure 3). The probes for chromosomes 12 and 21 both hybridize to one band in the size of ^~^1.8 Mb (Southern, bottom) matching the size expected if a chromosome fusion occurred and explaining the absence of their respective ‘original’ chromosomal bands.

**Figure S2:**
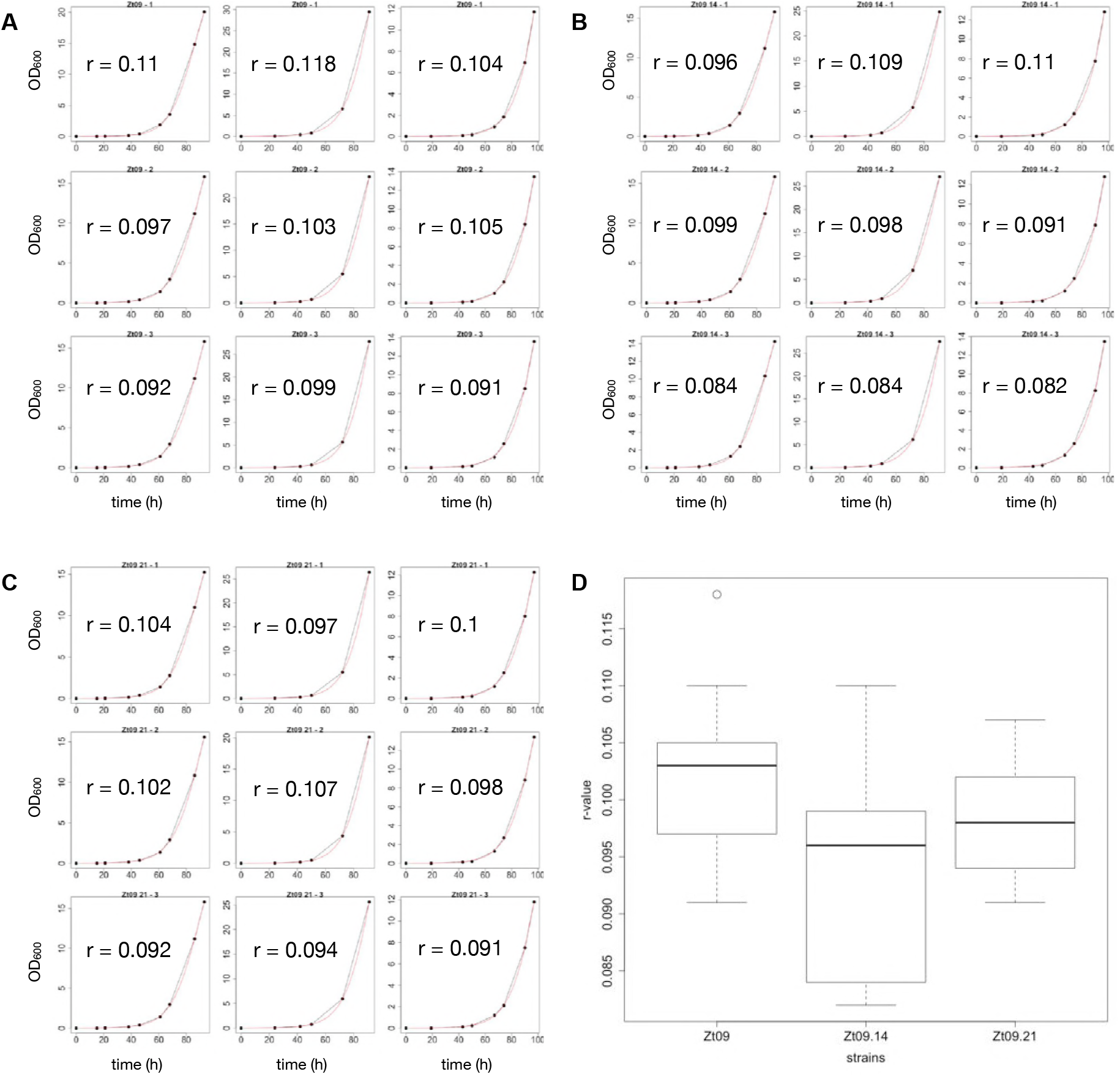
Growth assay of Zt09 and chromosome-loss strains Zt09Δ14 and Zt09Δ21. Three independent growth assays were conducted with the progenitor strain Zt09 (A) and two chromosome-loss strains Zt09Δ14 (B) and Zt09Δ21 (C), representing the loss of the largest and smallest accessory chromosome. Plotted are the fitted growth curves of all replicates and experiments generated with the R package growthcurver. The growth curves are based on OD_600_ measurements. Panel (D) displays a boxplot of all r values of the different strains. There are no significant differences between the growth curves of the three tested strains (Wilcoxon rank-sum test Zt09 – Zt09Δ14: p-value = 0.1443, Zt09 – Zt09Δ21: p-value = 0.3762).

**Figure S3:**
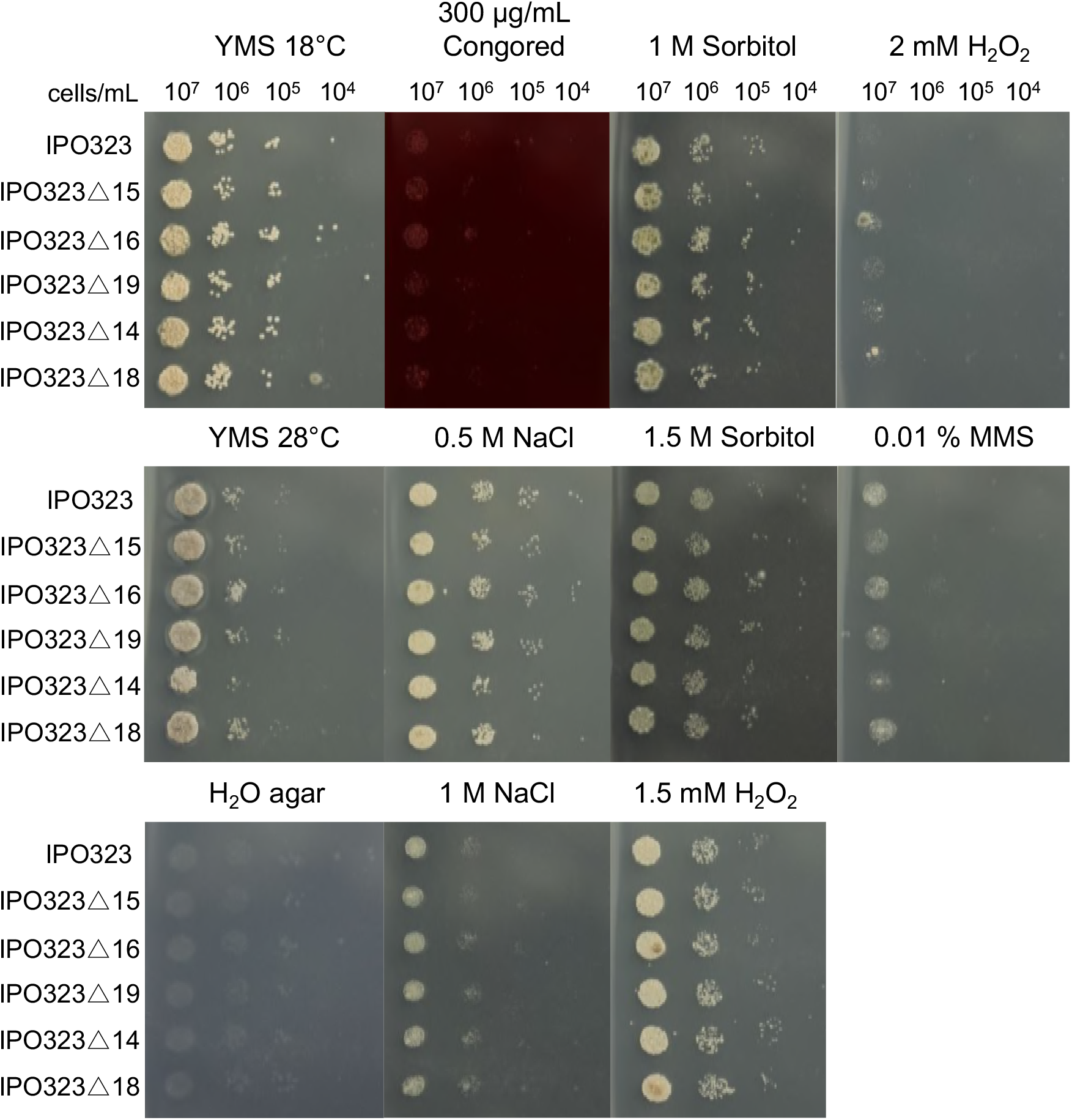
*In vitro* stress assays of the *Z. tritici* reference isolate IPO323 and the *in planta* chromosome-loss strains. Several *in vitro* stress conditions were tested to assess the effect of accessory chromosome losses on fitness. Five *in planta* chromosome-loss strains were tested, IPO323 was used as the reference strain. We observed no noteworthy differences in growth rate or colony morphology between reference and chromosome-loss strains.

## Supplementary text

Programs and commands used for quality filtering, mapping, visualization, and SNP calling for genome sequencing data and assembly of long-read SMRT sequencing reads of the *Zymoseptoria ardabiliae* reference strain Za17.

## Quality filtering

### Trimmomatic version 0.36 (Bolger et al. 2014)

java-jar trimmomatic-0.36.jar PE-threads 8-phred33 R1_sample_1.fastq.gz R2_sample_1.fastq.gz R1_paired.fastq R1_unpaired.fastq R2_paired.fastq R2_unpaired.fastq HEADCR0P:10 CROP:149 LEADING:3 TRAILING:3 SLIDINGWINDOW:4:15 MINLEN:50

## Mapping

### bowtie2 version 2.1.0 (Langmead & Salzberg 2012)

bowtie2-p8-q-x reference-1 R1_paired.fastq-2 R2_paired.fastq-S sample_1.sam

## Sorting and indexing

### samtools version 1.3.1 (Li 2011)

samtools view-Sb sample_1.sam > sample_1.bam

samtools sort sample_1.bam > sample_1_sorted.bam

samtools index sample_1_sorted.bam

## Extraction of discordant reads mapping to different chromosomes

samtools view-F 14-h input_sorted.bam | awk ‘($3!=$7 && $7!=“=“)’ > output_discordant.sam

## Visualization

### bedtools version 2.25.0 (Quinlan & Hall 2010)

bamToBed-i sample_1_sorted.bam > sample_1_sorted.bed

genomeCoverageBed-i sample_1_sorted.bed-g 2colums.chrom.sizes-bg > sample_1_sorted.bedGraph

## SNP calling

### samtools + bcftools version 1.3.1 (Li 2011)

samtools mpileup-E-C50-Q20-q20-uf reference.fasta sample_1_sorted.bam | bcftools call–ploidy 1-vc-O u-o sample_1.bcf

bcftools view sample_1.bcf > sample_1.vcf

bcftools filter-o sample_1_filtered.vcf-e’QUAL<20 | DP<10 | AF1<0.8’ sample_1.vcf

## Assembly of long-read SMRT sequencing *Zymoseptoria ardabiliae* reference strain Za17

High molecular weight DNA of the *Z. ardabiliae* strain Za17 was isolated as described in (Allen et al. 2006). Single molecule real time sequencing (SMRT) was performed on a PacBio RS II using 5 SMRT cells at the Functional Genomics Center Zürich, Switzerland.

The genome was assembled as previously described in (Plissonneau & Stürchler 2016) using the assembler SMRTanalysis v 2.3.0 implemented in the Hierarchical Genome Assembly Process version 3 (HGAP3) (Chin et al. 2013). Polishing was performed with Quiver that is part of the SMRTanalysis suite.

The following commands were used for the assembly:

> source Local_SMRTanalysis/current/etc/setup.sh
> fofnTOSmrtpipeInput.py HGAP.input.fofn > HGAP.input.xml
> smrtpipe.py-D NPROC=2-D MAX_THREADS=2-output=Result_SMRT – params=HGAP.input.xml xml:HGAP.input.xml

## Assembly statistics

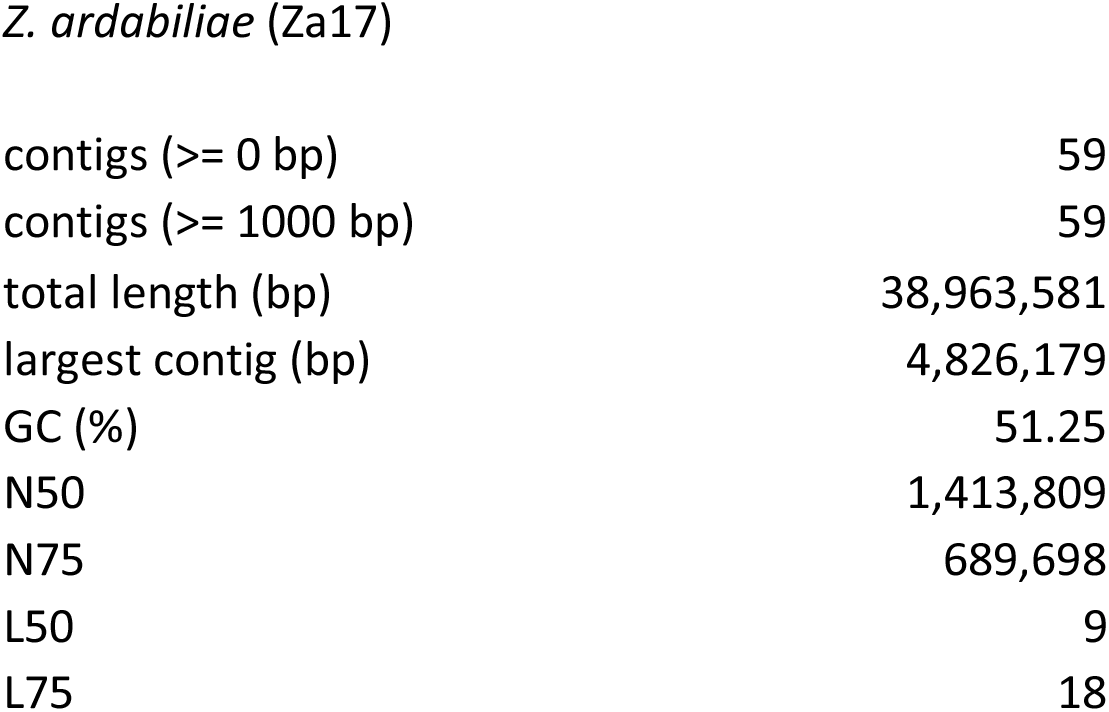

